# Comprehensive analysis of CXXX sequence space reveals that *S. cerevisiae* GGTase-I mainly relies on a_2_X substrate determinants

**DOI:** 10.1101/2024.03.04.583369

**Authors:** Anushka Sarkar, Emily R. Hildebrandt, Khushi V. Patel, Emily T. Mai, Sumil S. Shah, June H. Kim, Walter K. Schmidt

**Author notes:** **Communicating author:** Walter K. Schmidt, (email), 11.

## Abstract

Many proteins undergo a post-translational lipid attachment, which increases their hydrophobicity, thus strengthening their membrane association properties or aiding in protein interactions. Geranylgeranyltransferase-I (GGTase-I) is an enzyme involved in a three-step post-translational modification (PTM) pathway that attaches a 20-carbon lipid group called geranylgeranyl at the carboxy-terminal cysteine of proteins ending in a canonical CaaL motif (C - cysteine, a - aliphatic, L - often leucine, but can be phenylalanine, isoleucine, methionine, or valine). Genetic approaches involving two distinct reporters were employed in this study to assess *S. cerevisiae* GGTase-I specificity, for which limited data exists, towards all 8000 CXXX combinations. Orthogonal biochemical analyses and structure-based alignments were also performed to better understand the features required for optimal target interaction. These approaches indicate that yeast GGTase-I best modifies the Cxa[L/F/I/M/V] sequence that resembles but is not an exact match for the canonical CaaL motif. We also observed that minor modification of non-canonical sequences is possible. A consistent feature associated with well-modified sequences was the presence of a non-polar a_2_ residue and a hydrophobic terminal residue, which are features recognized by mammalian GGTase-I. These results thus support that mammalian and yeast GGTase-I exhibit considerable shared specificity.

**Article Summary:** This work investigates yeast GGTase-I specificity through genetics, high throughput sequencing, and two distinct reporter systems. This approach allows for comprehensive evaluation of all CXXX sequence space, which has not been possible with earlier approaches. We identified CXXX sequences supporting geranylgeranylation that differ from the historically defined CaaL sequence often cited in the literature as the GGTase-I target motif, and our results indicate that the last two amino acids of the target motif largely dictate GGTase-I specificity.

## Introduction

Protein lipidation involves the post-translational attachment of a lipid group to specific sites of a protein. These lipids can be fatty acids (palmitoyl, palmitoleyl myristoyl, octanoyl), isoprenoids (farnesyl, geranylgeranyl), sterols (cholesterol), and phospholipids (glycosylphosphatidylinositol) (Nadolski and Linder 2007, Resh 2013, Jiang, Zhang et al. 2018).

Protein prenylation is mediated by prenyltransferases - farnesyltransferase (FTase) that utilizes farnesyl pyrophosphate (FPP) as a substrate and three geranylgeranyltransferases (GGTase-I, -II, and -III) that utilize geranylgeranyl pyrophosphate (GGPP) (Benetka, Koranda et al. 2006, Wang and Casey 2016, Kuchay, Wang et al. 2019, Shirakawa, Goto-Ito et al. 2020). FTase and GGTase-I are considered CaaX-type prenyltransferases because their protein targets are defined by a C-terminal Ca_1_a_2_X motif: C - cysteine; a_1_, a_2_ – typically aliphatic amino acids (e.g., leucine, isoleucine, valine); X – many amino acids (**Figure 1A**). Where investigated, the X residue provides specificity for modification by FTase or GGTase-I (Moores, Schaber et al. 1991, Caplin, Hettich et al. 1994, Hartman, Hicks et al. 2005). FTase modifies a wide range of sequences where X is approximately half of the 20 amino acids, while GGTase-I modifies sequences where X is leucine, and sometimes phenylalanine, isoleucine, methionine or valine (i.e., Caa[L/F/I/M/V]) (Finegold, Johnson et al. 1991, Moores, Schaber et al. 1991, Yokoyama, McGeady et al. 1995, Hartman, Hicks et al. 2005, Maurer-Stroh and Eisenhaber 2005, Krzysiak, Aditya et al. 2010, Berger, Kim et al. 2018, Berger, Yeung et al. 2022, Kim, Hildebrandt et al. 2023). After CaaX protein prenylation, there is often but not always proteolytic removal of the -aaX residues by CaaX protease (Rce1) and carboxymethylation of the prenylated cysteine by isoprenylcysteine methyltransferase (ICMT). This multi-step modification is referred to in this study as canonical CaaX protein modification (**Figure 1A**). It is generally accepted that the PTMs of CaaX proteins augment their hydrophobic nature and are needed for their protein-membrane and protein-protein interactions (Maurer-Stroh, Washietl et al. 2003, Wang and Casey 2016). CaaX proteins serve a critical purpose in many cellular activities, such as signaling, growth, differentiation, and migration and relate to human diseases such as cancer, cardiovascular diseases, microbial infections, and progeria. Hence, CaaX-type prenyltransferases are often attractive targets for human disease therapies. (Benetka, Koranda et al. 2006, Berndt, Hamilton et al. 2011, Palsuledesai and Distefano 2015).

**Figure 1.**
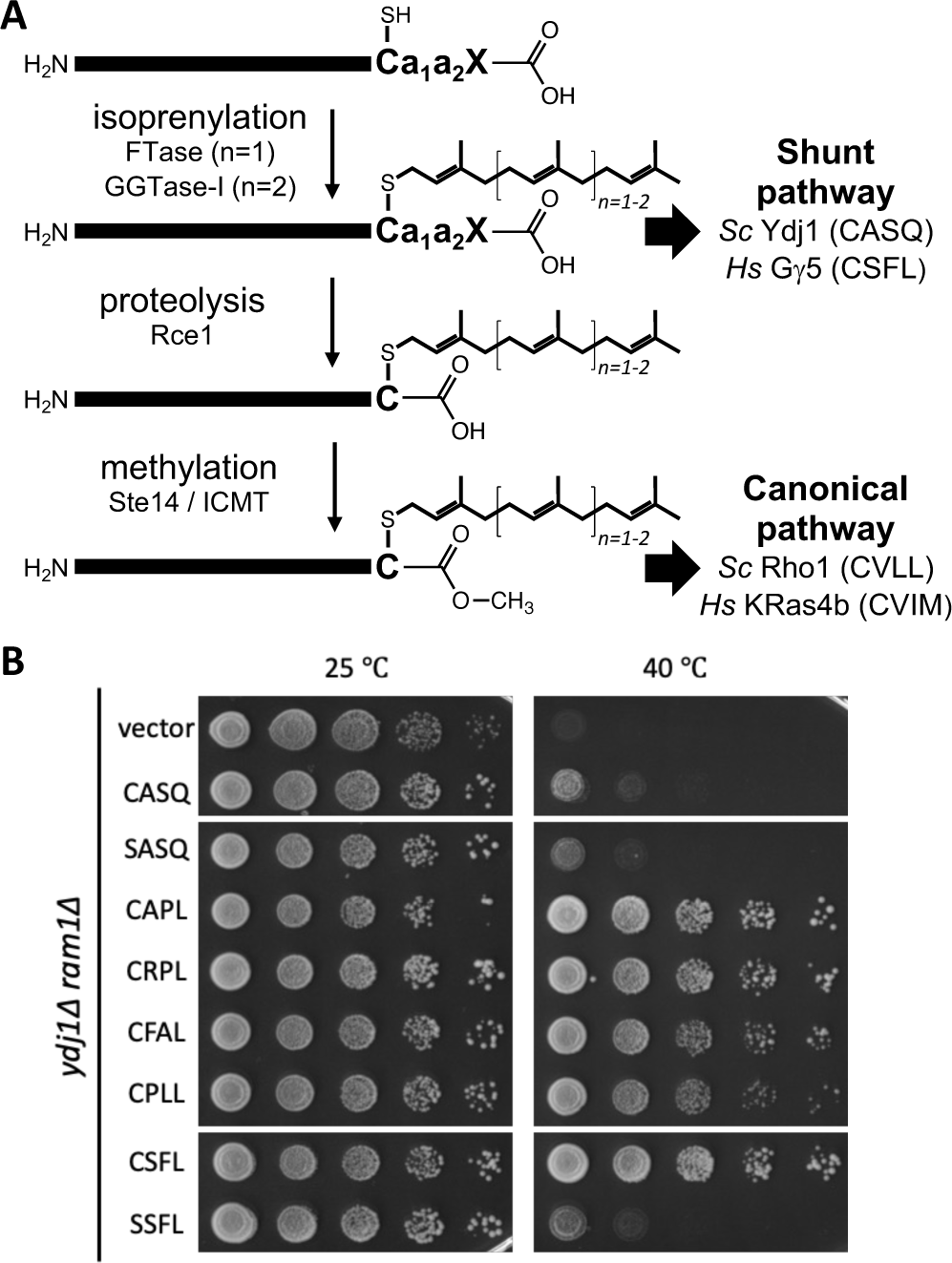
Geranylgeranylation can be investigated using a Ydj1 reporter. **A**) Overview of the CaaX protein modification pathway leading to either shunted (e.g., *Sc*Ydj1 or *Hs*Gγ5) or canonically modified proteins (e.g., *Sc*Rho1, *Hs*KRas4b). **B**) A Ydj1-based thermotolerance assay can be used to monitor GGTase-I activity. A yeast strain lacking Ydj1 and FTase (yWS2542, *ram1Δ ydj1Δ*) was transformed with plasmids encoding the indicated Ydj1-CXXX variants. Purified transformants were cultured to saturation in SC-Uracil, and a series of 10-fold dilutions were prepared and spotted onto YPD solid media. Plates were incubated at indicated temperatures for 96 hours (25 °C and 40 °C). Images are representative of results from experiments involving multiple biological and technical replicates.

Since the recognition of this three-step canonical CaaX protein PTM pathway, well-known prenyltransferase targets like the yeast **a**-factor (FTase), Ras GTPases (FTase), and Rho GTPases (GGTase-I) have led to the general view that the three steps of the pathway are coordinated. Emerging evidence reveals, however, that some CaaX proteins, like farnesylated yeast Ydj1 and geranylgeranylated mammalian Gγ5, can undergo prenylation but retain their last three amino acids (i.e., aaX) (Kilpatrick and Hildebrandt 2007, Hildebrandt, Cheng et al. 2016). This single-step prenylation-only modification is referred to in this study as “shunted” CaaX protein modification (**Figure 1A**). Shunted modification is required for optimal function of Ydj1, but the importance of shunting for other prenylproteins remains unexplored (Hildebrandt, Cheng et al. 2016).

Using the yeast Hsp40 Ydj1 protein as a genetic reporter, our studies have revealed that yeast FTase has much broader specificity than that defined by the canonical CaaX motif (Berger, Kim et al. 2018, Blanden, Suazo et al. 2018, Ashok, Hildebrandt et al. 2020, Kim, Hildebrandt et al. 2023). Many of the identified non-canonical sequences lack a_1_ and a_2_ branched-chain aliphatic amino acids and are not cleaved by Rce1, consistent with the observation that an aliphatic a_2_ residue is an important recognition determinant for Rce1 (Berger, Kim et al. 2018, Berger, Yeung et al. 2022, Kim, Hildebrandt et al. 2023). While the ability of mammalian FTase to modify non-canonical CaaX sequences has not been explored, farnesylated mammalian proteins with such sequences do exist, suggesting that shunted prenylproteins are widespread across species (e.g., DNAJA2, Lkb1/Stk11, Nap1) (Sapkota, Kieloch et al. 2001, Storck, Morales-Sanfrutos et al. 2019).

Studies utilizing *in vitro*, *in vivo* and *in silico* approaches have evaluated the specificity of FTase, but by comparison, the specificity of GGTase-I is underexplored. Several GGTase-I specificity studies have explored specificity using *in vitro* prenylation assays with purified GGTase-I and individually purified candidate proteins (Moores, Schaber et al. 1991, Caplin, Hettich et al. 1994). Medium throughput approaches have explored specificity using peptide libraries or metabolic probes to identify prenylated proteins within cells (Hartman, Hicks et al. 2005, Chan, Hart et al. 2009, Storck, Morales-Sanfrutos et al. 2019). None of these methods have allowed for evaluating GGTase-I against all possible 8000 CXXX sequence combinations. A prediction system for identifying GGTase-I targets has been established, but it too has limitations (Maurer-Stroh and Eisenhaber 2005). For example, the geranylgeranylated non-canonical sequence CSFL (Gγ5) is not predicted to be modified.

Overall, studies generally support that mammalian GGTase-I prefers CaaX sequences with aliphatic a_2_ and hydrophobic X residues. This specificity is entirely reliant on the β subunit of the dimeric GGTase-I complex, and conserved specificity among distinct GGTase-I enzymes is often observed (Caplin, Hettich et al. 1994, Mazur, Register et al. 1999, Reid, Terry et al. 2004, Benetka, Koranda et al. 2006). The specificity of yeast GGTase-I, however, has not been fully investigated. Rat and yeast GGTase-I β subunits have limited sequence conservation (27.2% identity; 39.9% similarity; 23.2% gaps per EMBOSS Needle), and active site residues that confer peptide substrate and lipid specificity are only partly conserved (**Table S1**) (Taylor, Reid et al. 2003, Reid, Terry et al. 2004).

Because yeast FTase has broader specificity than previously anticipated, and considering the low sequence conservation between mammalian and yeast GGTase-I, we have investigated the specificity of yeast GGTase-I. This was accomplished by genetic approaches that adapted the normally farnesylated yeast Hsp40 Ydj1 as a yeast GGTase-I reporter and used the established geranylgeranylated yeast GTPase Rho1 as a complementary reporter. Our results indicate that yeast GGTase-I targets the Cxa[L/F/I/M/V] sequence, where x is a wide range of residues, a is primarily one of the three branched-chain residues, and the terminal position is restricted to a limited set of nonpolar residues.

## Materials and Methods

### Yeast strains and plasmids

Yeast strains and plasmids used in this study are listed in **Table 1** and **Table S2**, respectively. Plasmid cloning strategies are described in **Table S3**.

**Table 1.** Yeast strains used in this study.

| Yeast strain | Genotype | Source |
| --- | --- | --- |
| yWS2542 | *MATa his3Δ1 leu2Δ0 met15Δ0 ura3Δ0 ram1::KAN ydj1::NAT* | (Berger, Kim et al. 2018) |
| BY4741 | *MATa his3Δ1 leu2Δ0 met15Δ0 ura3Δ0* | (Brachmann, Davies et al. 1998) |
| yWS3204 | *MATα his3Δ1 leu2Δ0 ura3Δ0 ram1::KAN* | This study |
| yWS1632 | *MATa his3Δ1 leu2Δ0 ura3Δ0 met15Δ0 ram1::KAN* | (Giaever, Chu et al. 2002) |
| ypr165w | *MATa/α his3Δ1/his3Δ1 leu2Δ0/leu2Δ0 ura3Δ0/ura3Δ0 MET15/met15Δ0 LYS2/lys2Δ0 RHO1/rho1::KAN* | Dharmacon |
| yWS3275 | *MATa his3Δ1 leu2Δ0 lys2Δ0 met15Δ0 ura3Δ0 rho1::KAN [CEN URA3 RHO1]* | This study |
| yWS3761 | *MATa leu2 lys2Δ0 his3 ura3 rho1::KAN ram1::KAN [CEN URA3 RHO1]* | This study |
| yWS3169 | *MATa met15Δ0 his3Δ1 leu2Δ0 ura3Δ0 ram2::KAN ydj1::NAT [CEN HIS3 P<sub>PGK1</sub>-FNTA] [CEN LEU2 P<sub>PGK1</sub>-PGGT1B]* | (Hildebrandt, Sarkar et al. 2023) |
| yWS4277 | *MATa his3Δ1 leu2Δ0 ura3Δ0 ram1::KAN ydj1::NAT [CEN HIS3 P<sub>PGK1</sub>-RAM2] [CEN LEU2 P<sub>PGK1</sub>-CDC43]* | This study |

yWS3761 was constructed using standard yeast genetic techniques involving a cross between *rho1* [*RHO1*] and *ram1*1 haploid strains and subsequent phenotypic and PCR analysis of haploid candidates arising from random sporulation of the diploid. The diploid precursor to yWS3761 was transformed with pWS1807 (*2µ LEU2 RAM1*) to facilitate sporulation, which was not otherwise evident. The *rho1*1 parent haploid strain yWS3275 was constructed from a commercially available heterozygous diploid strain ypr165w that was transformed with pWS1835 (*CEN URA3 RHO1*) and subjected to sporulation and random spore analysis to identify haploid *rho1*1 [*RHO1*] candidates with desired genetic markers. The chromosomal deletion of *RHO1* was confirmed by PCR and sensitivity to FOA. The *ram1* parent haploid strain yWS3204, which is a *MAT*α derivative of yWS1632 (Giaever, Chu et al. 2002), was constructed by several backcrosses to isogenic wildtype parent strain BY4742 to eliminate a petite phenotype. The chromosomal deletion of *RAM1* was tracked by G418 resistance.

Unless described otherwise, plasmids encoding Ydj1-CXXX and HA-tagged Rho1-CXXX variants were constructed by recombination-mediated PCR-directed plasmid construction in yeast (Oldenburg, Vo et al. 1997). Oligonucleotides (IDT, Newark, NJ) encoding for different CXXX variants were PCR amplified and co-transformed into yeast (BY4741) along with *NheI* / *AflII* linearized pWS1132 for the Ydj1-CXXX plasmids and *BamHI / BstAPI* linearized pWS2125 for the HA-tagged Rho1-CXXX plasmids. Colonies with recombinant plasmids were selected on SC-Uracil or SC-Leucine, followed by the isolation of plasmids from individual colonies and their verification by diagnostic restriction digests and sequencing of plasmid DNAs (Azenta, Burlington, MA). Construction of the Ydj1-CXXX encoded plasmids obtained from the Ydj1-CXXX (pWS1775) library has been described previously (Kim, Hildebrandt et al. 2023).

pWS1807 was constructed in two steps. First, a PCR product encoding the *RAM1* genomic locus was amplified from BY4741 that was co-transformed into yeast BY4741 with *HindIII* linearized pRS316 to allow for recombination-mediated PCR-directed plasmid construction *in vivo* (Oldenburg, Vo et al. 1997). The *SacI*-*XhoI* fragment encoding *RAM1* from the resultant plasmid pWS1767 was then subcloned into pRS425 at the same sites to create pWS1807. Diagnostic restriction digests and DNA sequencing (Azenta, Burlington, MA) were used to identify candidates at each step.

### Thermotolerance assay

This assay was performed as described previously (Hildebrandt, Cheng et al. 2016, Berger, Kim et al. 2018, Blanden, Suazo et al. 2018, Ashok, Hildebrandt et al. 2020, Kim, Hildebrandt et al. 2023). Briefly, yeast cells were cultured until saturation in SC-Uracil liquid media at 25 °C, and a portion of the saturated culture (100 µl) was added to the first well of a 96-well plate. A series of 10-fold dilutions were prepared and the dilution series spotted onto YPD plates. These plates were incubated at 25 °C and 40 °C for 96 hours and imaged with a Cannon flat-bed scanner (300 dpi; TIFF format). Photoshop was used for minor adjustments to images (i.e., contrast, rotation, cropping). This assay was performed twice on separate days with at least two technical replicates included in each trial.

### Ydj1-based thermotolerance screen

The *E. coli* derived Ydj1-CXXX plasmid library (pWS1775) has been previously described (Kim, Hildebrandt et al. 2023). This library contains all 8000 Ydj1-CXXX variants. The library was transformed into yWS2542 (*ram11 ydj11*) using the Frozen-EZ Yeast Transformation II Kit (Zymo Research, Irvine, CA) per the manufacturer’s guidelines. Approximately 367,000 colonies were collected by scraping cells off multiple SC-Uracil plates and washing them in SC-Uracil liquid media. The resuspended solution was centrifuged to obtain a cell pellet that was resuspended in 15% glycerol solution and stored at -80 °C as aliquots.

For each replicate of the thermotolerance screen, ∼150,000 CFUs were used. This number of CFUs ensured >99.9% coverage of the Ydj1-CXXX library (http://guinevere.otago.ac.nz/cgi-bin/aef/glue.pl). Briefly, ∼55 x 10^6^ cells were inoculated into 200 mL SC-Uracil liquid media in duplicate, each mixture split into 10 x 20 mL replicates, and each 10-member set subsequently incubated at permissive temperature (25 °C) or restrictive temperatures (37 °C and 42 °C) for 24 - 48 hours until saturation (A_600_ 1.9 – 3.3). This growth period was estimated to allow the cells to go through at least 8 rounds of population doubling. Cells from the saturated cultures were harvested by centrifugation, washed, and collected. Plasmids were extracted from the cells using E.Z.N.A. Yeast Miniprep kit following manufacturer’s guidelines (OMEGA Bio-Tek, Norcross, GA). Of note, the high temperatures used for liquid-based growth were 37 °C and 42 °C, while 40 °C was chosen for plate-based growth.

### Next-Generation sequencing

The library preparation for Next-Generation Sequencing (NGS) (i.e., Ydj1-CXXX plasmid libraries derived from the thermotolerance screens performed at 25 °C, 37 °C, and 42 °C) and the NGS method itself have been described previously (Kim, Hildebrandt et al. 2023). In addition, ten replicates of the *E. coli* derived and the naïve yeast derived Ydj1-CXXX plasmid libraries were sequenced by NGS. The NGS run resulted in over 6 million reads with 97% of read having a Q30 quality score or better (i.e., 99.9+% base call accuracy). The frequency of each CXXX sequence within each library (i.e., *E. coli* derived, naïve yeast derived, 25 °C, 37 °C and 42 °C) was calculated by summing the occurrence of a specific CXXX sequence in all ten replicates and dividing that value by the sum of all CXXX sequence occurrences in the data set (**File S1**). Ultimately, each CXXX sequence had two NGS Enrichment Scores (NGS E-Score), which was the frequency of a specific CXXX sequence at the restrictive temperature (37 °C or 42 °C) divided by its frequency in the naïve yeast library (**File S1**).

### YDJ1-CXXX-based mini-screen to sample CXXX sequences

Ydj1-CXXX library (pWS1775) plasmid library was transformed into yWS2542 (*ram11 ydj11*) and incubated on SC-Uracil media plates at 25 °C. Individual transformants (i.e., colonies) were cultured in preparation for the thermotolerance assay at 25 °C and 40 °C. Briefly, cultures of the individual transformants were subjected to a fixed dilution, and the single dilution spotted on YPD media plates that were incubated at 25 °C and 40 °C. A collection of 20 candidates displaying thermotolerant and thermosensitive phenotypes at 40 °C was identified, and plasmids extracted from each candidate were recovered and sequenced by Sanger sequencing. The collection was reduced to 8 CXXX sequences that were judged to best represent an even distribution over the complete range of the NGS E-Score plots.

### Ydj1-based gel-shift assay

This assay was done as described previously (Berger, Kim et al. 2018, Hildebrandt, Sarkar et al. 2023). Briefly, yeast strains were cultured in SC-Uracil liquid media to log or late log phase (A_600_ 0.95 - 1.45) and total cell lysates were prepared by alkaline hydrolysis and trichloroacetic acid precipitation (Kim, Lapham et al. 2005). Total cell lysates were subjected to SDS-PAGE (9.5% separating gel) followed by immunoblot. The primary antibody used was rabbit anti-Ydj1 (courtesy of Dr. Avrom Caplan) and the secondary antibodies were HRP-conjugated donkey or goat anti-rabbit antibodies (Kindle Biosciences, LLC). The blots were treated with ECL spray (ProSignal Pico Spray) and digital images captured using KwikQuant Imager system (Kindle Biosciences). Photoshop was used for minor adjustments to images (i.e., contrast, rotation, cropping). This assay was performed with two biological replicates for the yWS2542 and yWS3169 transformants and one biological replicate for the yWS4277 transformants.

### Rho1-based plasmid loss assay

yWS3761 (*rho1Δ ram1Δ [CEN URA3 RHO1])* was transformed with *LEU2* marked HA tagged Rho1-CXXX variants and grown on SC-Uracil and Leucine solid media (SC-UL) at 25 °C. The purified transformants were inoculated in SC-Leucine liquid media and cultured to saturation at 25 °C. The saturated cultures were diluted to 1 A_600_, and 100 μL of each diluted culture was transferred to an independent well of a 96-well plate, which was used to prepare a 10-fold serial dilution for each sample. The dilution series were collectively pinned onto YPD as a control for preparation of the dilution series and SC complete media with 5-FOA for the functional assay. The plates were incubated at 25 °C for 72 hours and scanned with a Cannon flat-bed scanner (300 dpi; TIFF format). Photoshop was used for minor adjustments to images (i.e., contrast, rotation, cropping). This assay was performed twice on separate days with at least two technical replicates included in each trial.

### Rho1-based cell viability selection

yWS3761 (*rho1Δ ram1Δ [CEN URA3 RHO1])* was co-transformed with *BamHI* linearized pWS2125 and PCR amplified CXXX mutants. The transformation reactions were plated onto SC-Leucine solid media and incubated at 25 °C. Colonies surviving selection were replica plated onto SC complete media with 5-fluoroorotic acid (5-FOA), and 200 FOA-resistant single colonies were individually isolated. Each candidate was patched onto 5-FOA media to confirm the 5-FOA resistant phenotype, then replica plated onto SC-Leucine to confirm leucine prototrophy, indicative of a *CEN LEU2 HA-RHO1-CXXX* plasmid being present. Candidates with all requisite phenotypes were cultured in SC-Leucine liquid media, and their associated plasmids recovered and sequenced (Azenta, Burlington, MA). Each plasmid was then individually re-transformed into yWS3761, and the Rho1-based plasmid loss assay was repeated with the transformant to confirm the FOA-resistant growth phenotype imparted by the plasmid.

### WebLogo Analysis

This was done as previously described with certain modifications (Berger, Kim et al. 2018). The desired groups of sequences were analyzed using the Weblogo website server (https://weblogo.berkeley.edu/logo.cgi) by amino-acid frequency-based analysis (Crooks, Hon et al. 2004). A customized amino acid coloring scheme was used: Cys was denoted blue, branched-chain aliphatic amino acids (Val, Ile, and Leu) were denoted red, charged amino acids (Lys, Arg, His, Asp, Glu) were denoted green, polar and uncharged residues (Ser, Gln, Thr, Asn) were denoted black and all other hydrophobic amino acids (Ala, Gly, Pro, Phe, Trp, Tyr and Met) were denoted purple.

## Results

### Ydj1 can be used as an efficient genetic reporter for yeast GGTase-I activity

Growth of yeast at elevated temperature requires prenylation of the Ydj1 Hsp40 chaperone, but not the downstream protease and carboxylmethylation steps of the CaaX modification pathway (Hildebrandt, Cheng et al. 2016). Due to this property, Ydj1 is a more direct reporter for prenylation. This Ydj1-dependent phenotype has been previously used for probing yeast FTase specificity (Berger, Kim et al. 2018, Kim, Hildebrandt et al. 2023).

Using Ydj1 as a reporter has led to the discovery of many farnesylatable CXXX sequences that do not adhere to the canonical CaaX consensus (Berger, Kim et al. 2018, Kim, Hildebrandt et al. 2023). A lot of these sequences support a robust Ydj1-dependent thermotolerance phenotype equivalent to that supported by wildtype farnesylated and shunted Ydj1 (CASQ). Within that study, a few CXX[L/F] sequences were determined to be prenylated in the absence of FTase activity, ostensibly by GGTase-I. To extend this observation and explore the potential utility of Ydj1 as an efficient genetic reporter for GGTase-I activity, we performed additional thermotolerance assays using the FTase deficient strain (i.e., *ram1*Δ) and certain selected Ydj1-CXXL sequences. This study confirmed that neither wildtype Ydj1 (CASQ) nor an unmodifiable Ydj1 variant (SASQ) can support growth at high temperature (40 °C) (**Figure 1B**). Yet, Ydj1 was able to support thermotolerant growth in the context of CAPL, CRPL, CFAL, CPLL and CSFL sequences. These sequences all contain Leu at the X position but lack a branched-chain aliphatic amino acid at the a_1_ position of the CaaX motif, and in all but one case, also lack such an amino acid at the a_2_ position. Moreover, Ydj1-CSFL supported a robust thermotolerant phenotype by comparison to unmodified Ydj1-SSFL, indicating that thermotolerance in the FTase-deficient background was cysteine-dependent. CSFL is associated with mammalian Gγ5, which is the only reported example of a shunted geranylgeranylated sequence; it lacks a_1_ and a_2_ amino acids needed for recognition and cleavage by the Rce1 CaaX protease (Trueblood, Boyartchuk et al. 2000, Kilpatrick and Hildebrandt 2007, Berger, Kim et al. 2018, Berger, Yeung et al. 2022). Together, these observations indicate that yeast Ydj1 can be utilized as a reporter for GGTase-I activity. Additionally, the data suggest that GGTase-I specificity is broader than the canonical Caa[L/F/I/M/V] consensus, especially with respect to requiring branched-chain aliphatic amino acids at the a_1_ and a_2_ positions.

### Canonical and non-canonical CaaX sequences impart different effects in the Ydj1-based thermotolerance assay

To investigate the possibility that yeast GGTase-I has broader specificity than anticipated, we designed a study to examine the ability of GGTase-I to modify all 8000 possible CXXX variants. We adapted a high throughput Ydj1-based thermotolerance screen that had been used previously by *Kim et. al*, 2023 to evaluate yeast FTase specificity (**Figure 2A**). In our case, we evaluated thermotolerance using a strain lacking FTase so that any identified thermotolerant phenotypes could be specifically attributed to GGTase-I activity.

**Figure 2.**
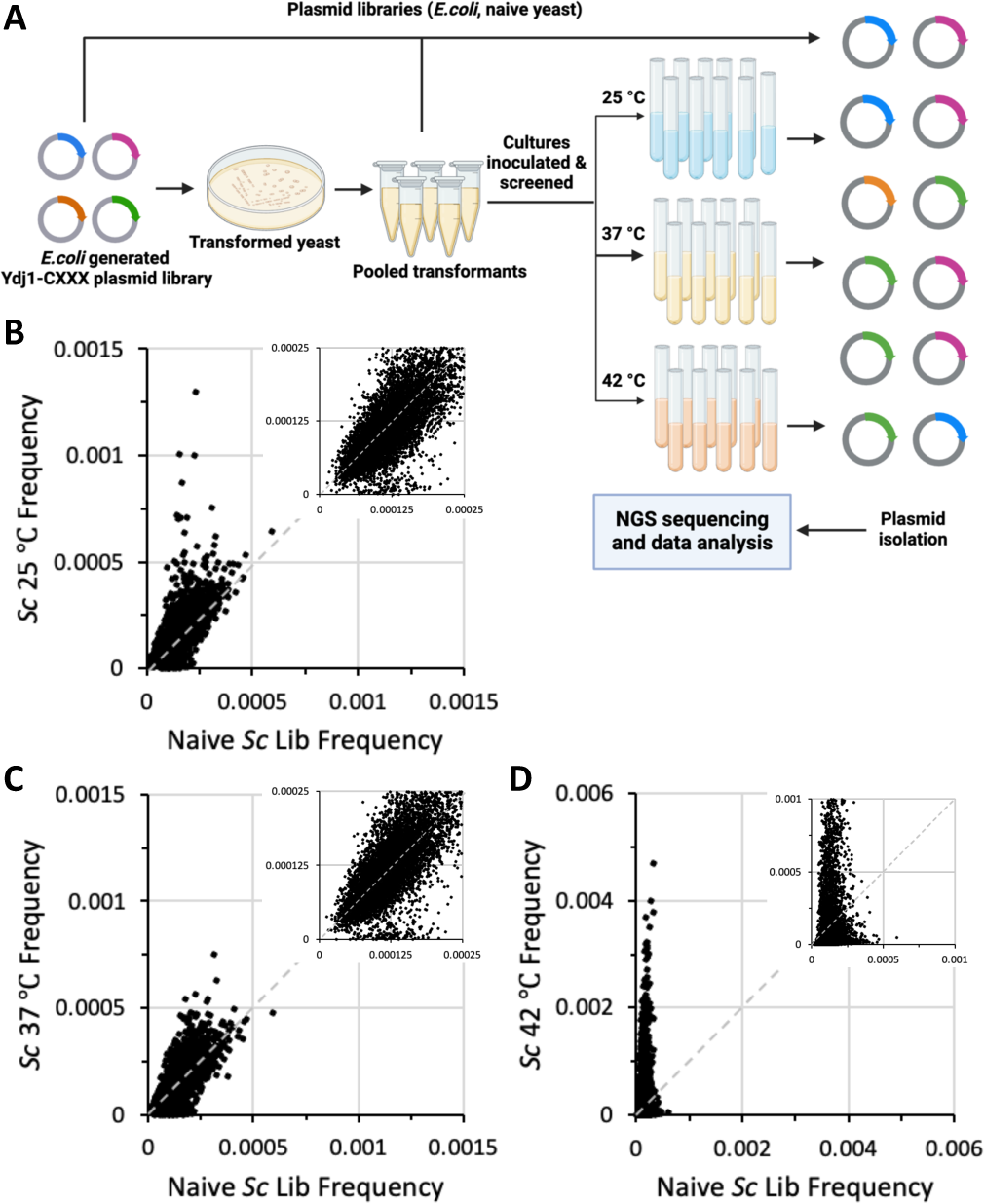
A thermotolerance screen yields an enriched population of Ydj1-CXXX variants. **A**) Experimental strategy for probing the entirety of CXXX space that can be modified by yeast GGTase-I. A plasmid library containing all 8000 Ydj1-CXXX combinations was created and transformed into a yeast strain lacking Ydj1 and FTase (yWS2542, *ram1Δ ydj1Δ*). The transformed colonies were harvested from multiple plates, pooled, and a representative aliquot used to inoculate cultures that were incubated at permissive and restrictive temperatures (25 °C, 37 °C and 42 °C, respectively) until saturation. Plasmids isolated from all populations were sequenced using high throughput methods, and data analyzed to determine the relative frequency of each Ydj1-CXXX variant in each population. Graphic created using BioRender.com. **B-D**) Plots of frequencies of sequences observed in naïve yeast library relative to those observed after cell expansion under B) 25 °C, C) 37 °C, or D) 42 °C conditions. Inset is a magnification of points near the origin.

Several large data sets were created as part of this study. The *E. coli* derived Ydj1-CXXX plasmid library has been previously characterized and is known to contain all 8000 Ydj1-CXXX variants (Kim, Hildebrandt et al. 2023). This *E. coli* derived library was transformed into *ydj1*1 *ram1*1 yeast, and colonies were recovered directly from the transformation plates to create a naïve yeast plasmid library. This yeast-derived library was confirmed by NGS to also contain all 8000 Ydj1-CXXX variants. Comparison of the frequencies for each CXXX sequence within the *E. coli* and naïve yeast libraries revealed no obvious enrichment bias for any Ydj1-CXXX variants due to the transfer into yeast (**Figure S1**). The naïve yeast library was inoculated into liquid media and propagated under both permissive (25 °C) and restrictive (37 °C and 42 °C) temperatures, where the restrictive temperatures were expected to enrich for cells expressing geranylgeranylated Ydj1-CXXX variants. NGS was utilized to evaluate the frequency of Ydj1-CXXX variants in all the expanded yeast populations (**Figure 2B-D**). Although we expected to observe no significant difference in CXXX frequencies between the naïve yeast and 25 °C libraries, we observed both enriched and de-enriched outliers in the 25 °C sample (**Figure 2B**). Comparison of CXXX frequencies in the naïve yeast and elevated temperature libraries (37 °C and 42 °C) revealed both enriched and de-enriched CXXX motifs in the 37 °C sample and strong enrichment of many CXXX motifs in the 42 °C sample (**Figure 2C and 2D**). To determine the relative enrichment of each Ydj1-CXXX variant at the restrictive temperatures, an enrichment score (NGS E–Score) was calculated for each CXXX sequence by dividing the frequency of a specific sequence in the restrictive temperature (37 °C or 42 °C) library relative to its frequency in the naïve yeast library. Those with higher NGS E-Scores were deemed as having better growth than others at the restrictive temperature. This method of analysis was previously used to interrogate FTase specificity using Ras and Ydj1 reporters (Stein, Kubala et al. 2015, Kim, Hildebrandt et al. 2023). While those previous studies used frequency data from 25 °C samples as denominators for deriving NGS E-Scores, we rejected this approach due to the observation of outliers in the 25 °C sample, which could result in false negative and positive outcomes.

2D plots revealed NGS E-Scores ranging from 0 – 3.2 (37 °C vs. naïve yeast) and 0 – 21.2 (42 °C vs. naïve yeast), indicative that enrichment (>1) and de-enrichment (<1) of Ydj1-CXXX variants had indeed occurred (**Figure 3A and 3B**). The sequences evaluated in preliminary studies were also observed to be broadly distributed in these 2D plots. Ydj1-CASQ was de-enriched and in the bottom 25% of hits in the 42 °C library, which was consistent with its expected lack of geranylgeranylation. At 37 °C, which was a less stringent condition, CASQ was neither enriched nor de-enriched. Ydj1 harboring non-canonical sequences CRPL, CPLL, CFAL, CAPL and CSFL expected to be enriched (see **Figure 1**) were present in the top 15% hits in both the 37 °C and 42 °C libraries, which was consistent with their expected geranylgeranylation. Notably, CXXX sequences associated with yeast (*Sc*) Rho GTPases (CVLL, CIIL, CTIM, CIIM, CVIL, CAIL), *Sc* Ras-related GTPase Rsr1 (CTIL) and human (*Hs*) K-Ras4b (CVIM) were among the de-enriched population in both the libraries. These sequences are frequently touted as examples of canonical geranylgeranylated or cross-prenylated sequences (i.e., modified by both FTase and GGTase-I) (Caplin, Hettich et al. 1994). While not initially predicted, the de-enrichment of these sequences concurs with observations that farnesylated Ydj1 that is canonically modified has impaired activity (i.e., Ydj1-CVIA), although such modified sequences still outperform unmodified Ydj1 (i.e., Ydj1-SASQ) (Hildebrandt, Cheng et al. 2016). In this study, by contrast, Ydj1-CXXX variants predicted to be geranylgeranylated and canonically modified vastly underperform unmodified Ydj1 (i.e., Ydj1-CASQ) under the competitive growth conditions of the thermotolerance screen. We were able to recapitulate these differences for individual Ydj1-CaaX variants using a plate-based assay (**Figure S2**). Taken together, these observations indicate that the Ydj1-based thermotolerance screen enriches for shunted CXXX motifs relative to unmodified and canonically modified sequences. Additionally, these observations suggest that an increased hydrophobicity imparted by more hydrophobic geranylgeranyl (vs. farnesyl) and carboxylmethylation (vs. free carboxyl) can synergistically impair Ydj1 activity.

**Figure 3.**
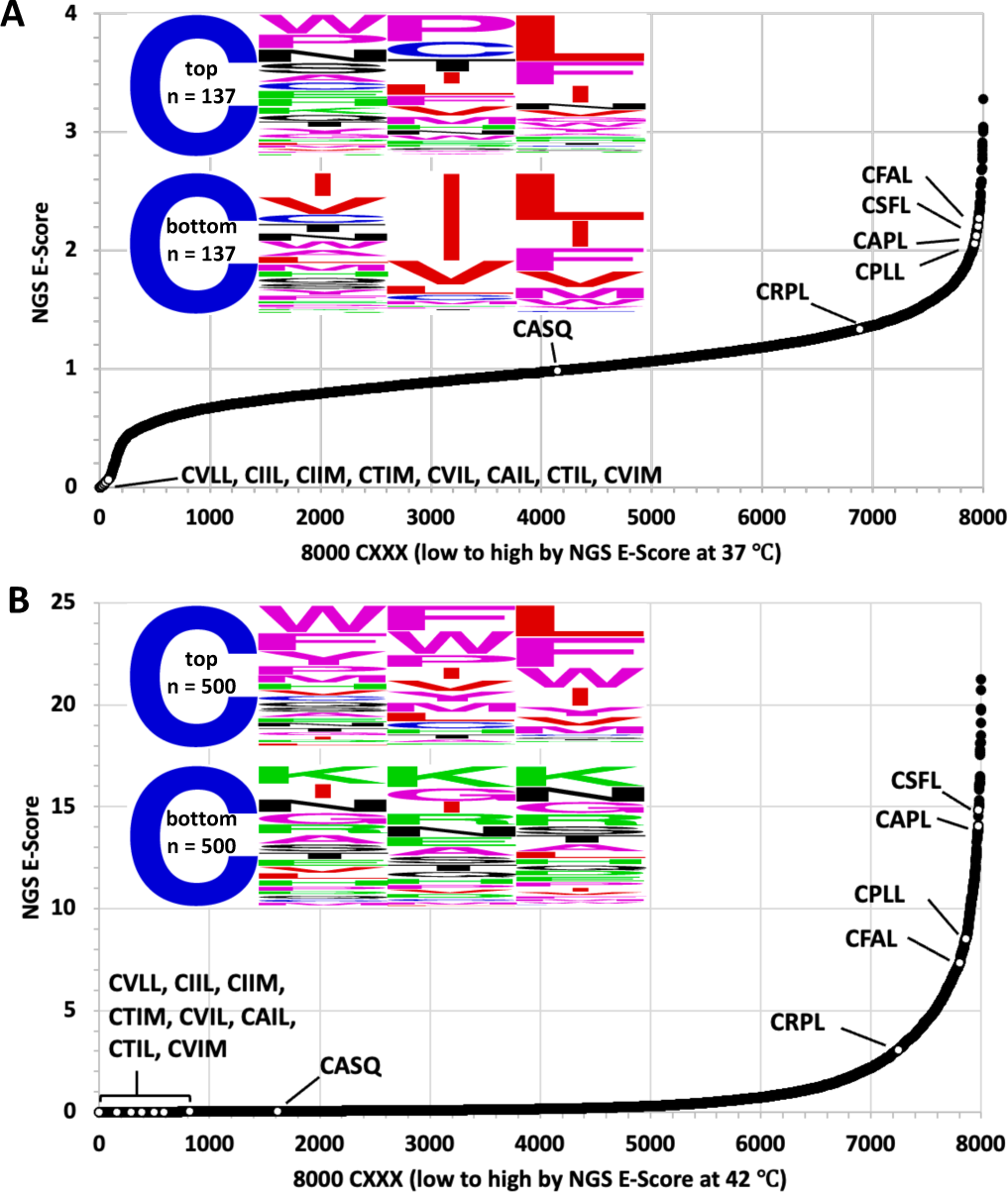
Temperature effects on the *enrichment and de-enrichment of Ydj1-CXXX variants.* Enrichment scores (NGS E-Score) for each of the 8000 Ydj1-CXXX sequences were determined relative to the naïve yeast library for sequences in the **A**) 37 °C and **B**) 42 °C yeast libraries. The NGS E-Scores are represented as 2D plots with white dots representing the scores of different sequence sets: *i*) proteins known or highly suspected to be geranylgeranylated, or not geranylgeranylated (i.e., controls) – *Sc* Rho1 (CVLL), *Sc* Rho2 (CIIL), *Sc* Rho3 (CIIM), *Sc* Rho4 (CTIM), *Sc* Rho5 (CVIL), *Sc* Cdc42 (CAIL), *Sc* Rsr1 (CTIL), *Hs* K-Ras4b (CVIM), *Sc* Ydj1 (CASQ), *ii*) sequences from the thermotolerance pilot study described in Figure 1B (CRPL, CFAL, CPLL, CAPL), and *iii*) the sequence derived from *Hs* Gγ5 (CSFL). The WebLogos associated with each plot reflect an analysis for a subset of sequences. In panel A, the analysis was performed using the 137 highest NGS E-Scores associated with the 37 °C data set (top), and an equivalent number of sequences with the lowest NGS E-Scores <0.2 (bottom). In panel B, the analysis was performed using sequences reflecting the top 500 NGS E-Scores associated with the 42 °C data set (top), and equivalent number of sequences with the lowest NGS E-Scores (bottom).

A comparison of the 2D plots indicated a striking difference in the curve profiles. Significant de-enrichment was observed for some sequences in the 37 °C data set that was not observed in the 42 °C data set (i.e., compare left most regions of 2D plots). WebLogos were created to better visualize the amino acids associated with this region in both plots, as well as the corresponding enriched regions. Highly de-enriched sequences associated with the 37 °C data (NGS E-Scores 37 °C / naïve *Sc* library < 0.2) captured all the Rho/Ras-related sequences clustered in this region of the 2D plot. A Weblogo of this subset (n=137) had a near canonical-looking profile Cx[V/I/L][L/F/I/M/V]. The dominance of branched-chain aliphatic residues at the a_2_ position for these sequences, which are optimal for Rce1 CaaX protease specificity, indicate that this region of the 2D plot appears to be populated mainly by canonical geranylgeranylated sequences (**Figure 3A**, bottom panel). By comparison, the equivalent number of top performing sequences in this data set (n=137) contained a wider range of a_2_ amino acids and more variability at the X position (**Figure 3A**, top panel).

A similar WebLogo analysis was performed using the 42 °C data set. A larger subset of de-enriched sequences (n=500) was analyzed to capture most, albeit not all, of the Rho/Ras-related sequences clustered in this region (**Figure 3B**, bottom panel). The sequence profile revealed a wide range of amino acids, including residues at a_2_ (i.e., D/E/K/R) or X (i.e., K/R/P) that interfere with farnesylation of Ydj1, which we propose are also likely to interfere with geranylgeranylation (Berger, Kim et al. 2018, Kim, Hildebrandt et al. 2023). In fact, over half of the de-enriched sequences contained these restrictive amino acids (**Figure S3A**). A smaller subset of sequences matching the canonical GGTase-I consensus were also among the de-enriched population (**Figure S3B**). The remaining sequences in the population displayed no obvious pattern (**Figure S3C**). A WebLogo analysis of the most enriched sequences in this data set was also performed (**Figure 3B**, top panel). This analysis revealed no obvious enrichment of aliphatic amino acids at a_1_ and a_2_ positions, but hydrophobic amino acids were generally enriched with aromatic amino acids F and W being most prevalent. A moderate enrichment of expected amino acids was observed at the X position (i.e., L/F/I/M/V), as were a few other amino acids (e.g., W and Y). Overall, this analysis indicates that the enriched sequences recovered by our screening approach do not fully adhere to the expected Caa[L/F/I/M/V] consensus sequence of GGTase-I.

### Enriched non-canonical CXXX sequences confer Ydj1-dependent thermotolerance

To validate the observations from the Ydj1-based thermotolerance screen through an orthogonal assay, we evaluated 15 CXXX sequences by Ydj1-based thermotolerance and gel-shift assays. Eight non-canonical CXXX sequences were derived from a Ydj1-based thermotolerance mini-screen (see Materials and Methods), and the others were CASQ (shunted farnesylated sequence of yeast Ydj1), CVLL and CVIL (canonical geranylgeranylated sequences of yeast Rho1 and Rho5, respectively), CSFL (shunted geranylgeranylated sequence of mammalian Gγ5), and CAFL, CPIQ and CHLF (sequences with high NGS E-Scores). These 15 candidates were chosen to represent broad distribution of scores across the NGS E-Score plots (37 °C vs. naïve yeast library and 42 °C vs. naïve *Sc* library) (**Figure 4A and 4B**).

**Figure 4.**
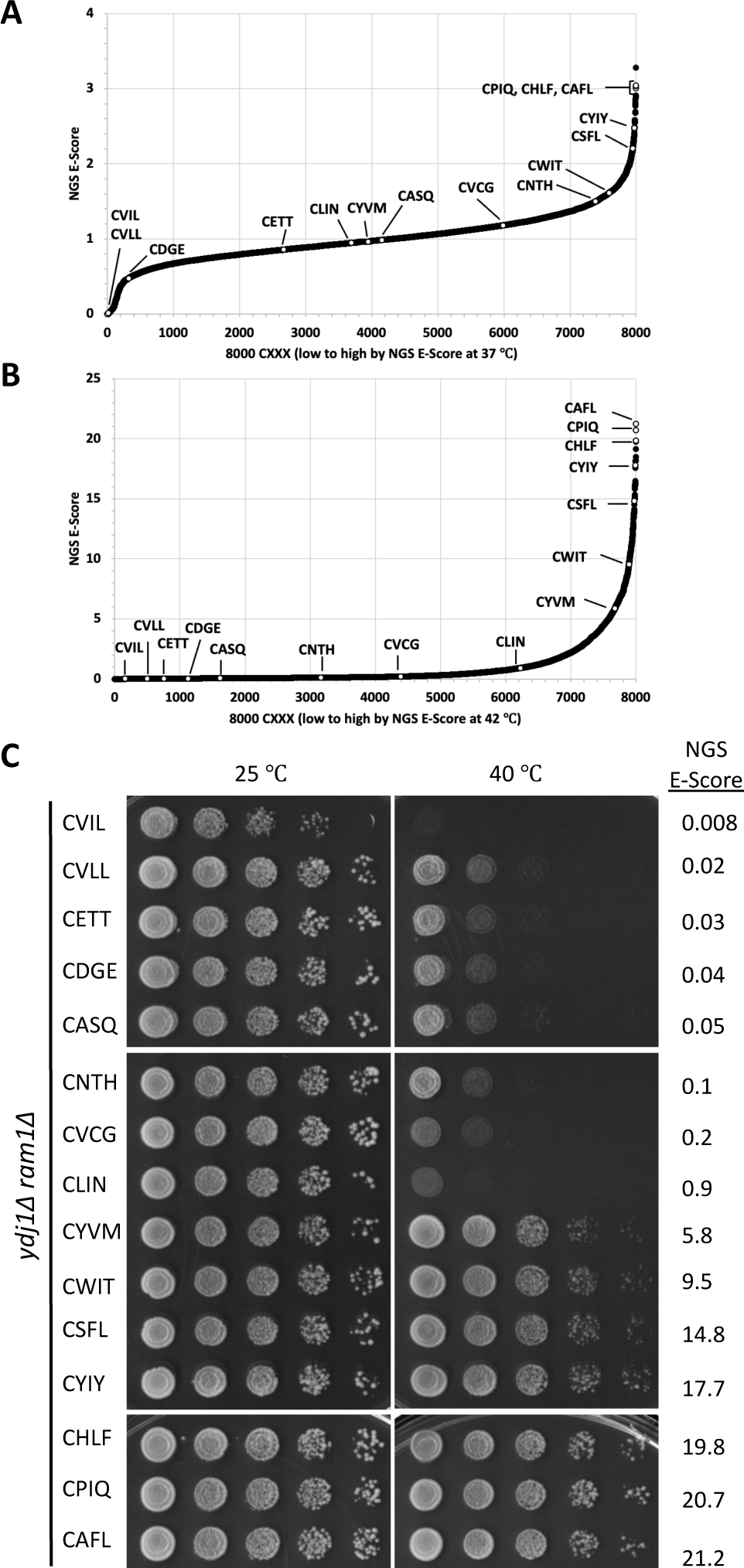
Validation of thermotolerance status for a representative set of Ydj1-CXXX variants. **A-B**) The white dots mark the position of 15 sequences that represent a wide distribution of NGS E-Scores on the plots described in **Figures 3A and 3B**. Included among the sequences representing the Test Set are several predicted to be geranylgeranylated (CSFL, CVLL, and CVIL) or not geranylgeranylated (CASQ) and three that had the highest NGS E-Scores from the 42 °C data set (CHLF, CPIQ, and CAFL). **C**) Thermotolerance assay results observed for the subset of sequences described in panel A. Sequences were evaluated as described in Figure 1B. NGS E-Scores refer to 42 ℃ library frequency vs. naïve yeast library frequency values.

The thermotolerance assay results indicated that 8 of 15 CXXX sequences could support robust growth at high temperatures. We generally observed consistency between NGS E-Score (42 °C vs. naïve *Sc* library) and high-temperature growth (**Figure 4C**). Higher E-scores had better thermotolerance. The thermotolerant phenotype appeared between NGS E-Scores of 0.9 and 5.8; a lack of data points between these NGS E-Scores did not allow for further refinement of a minimum threshold for thermotolerance. The canonical sequences CVIL and CVLL were among the 7 thermosensitive sequences.

### Enriched non-canonical CXXX sequences are subject to partial modification by yeast GGTase-I

In past studies of FTase specificity, a strong correlation has been observed between thermotolerance and farnesylation of Ydj1, where the latter was determined by gel-shift analysis (Berger, Kim et al. 2018). The basis for the gel-shift assay is that farnesylated Ydj1 migrates faster by SDS-PAGE than unfarnesylated Ydj1. We thus investigated whether this was also the case for Ydj1-CXXX variants that were predicted to be modified in the absence of FTase. Ydj1 harboring canonical sequences (i.e., CVLL and CVIL) exhibited a mobility shift relative to unmodified Ydj1p (i.e., CASQ) (**Figure 5A**). An obviously gel-shifted population was not observed for the other sequences evaluated, but we consistently observed a light smear beneath the main Ydj1 band for sequences having NGS E-Scores greater than or equal to 0.9 (i.e., CLIN, CYVM, CWIT, CSFL, CYIY, CHLF, CPIQ, and CAFL). To further investigate this observation, we performed the gel-shift assay with select CXXX sequences and their cysteine to serine mutants (SXXX) (**Figure 5B**). CVLL again exhibited faster migration and a gel-shift pattern that was clearly distinguishable from their unmodifiable serine counterpart. CASQ and CNTH, which lacked the very light smear, exhibited no obvious mobility difference relative to their serine counterparts. CLIN, CSFL, CPIQ and CAFL exhibited a very light smear beneath the main protein band, which was absent in the serine mutants. In some cases, cultures were incubated at 37 °C to try and improve gel-shift properties to no avail. Together, these observations suggest that many of these non-canonical geranylgeranylated sequences are modified by GGTase-I, albeit only partially, yet this partial modification is sufficient to impart a Ydj1-dependent thermotolerance phenotype. To confirm that sequences with higher NGS E-Scores were reactive with yeast GGTase-I, we overexpressed yeast GGTase-I in an effort to exaggerate geranylgeranylation. Indeed, we observed improved gel-shift patterns for sequences with higher NGS E-scores and no noticeable change for those with lower scores (**Figure 5C**). Of the modified sequences, none appeared to be fully modified.

**Figure 5.**
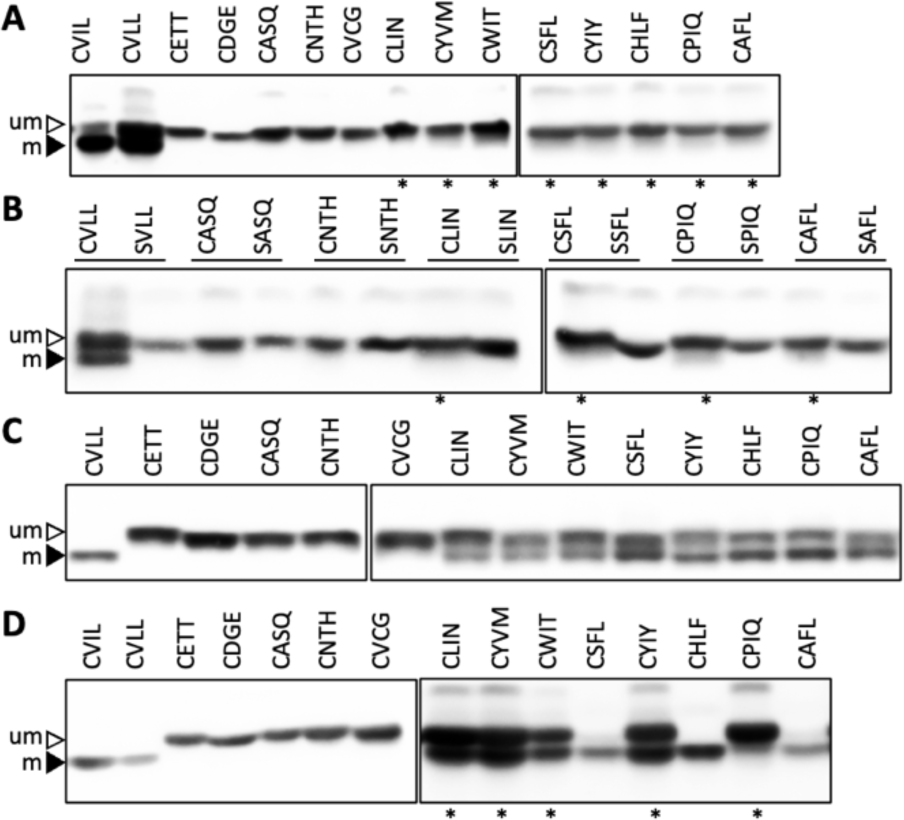
Evaluation of Ydj1-CXXX variants in the Test Set by gel-shift assay. A yeast strain lacking Ydj1 and FTase (yWS2542, *ram1Δ ydj1Δ*) was transformed with plasmids **A**) from the Test Set or **B**) matched pairs of sequences encoding either Ydj1-CXXX or its Ydj1-SXXX variant. **C-D**) The Ydj1-CXXX variants described in panel A were expressed in a yeast strain lacking Ydj1 that overexpresses either C) yeast GGTase-I (yWS4277, *ram1Δ ydj1Δ [CEN HIS3 P_PGK_-RAM2] [CEN LEU2 P_PGK_-CDC43]*) or D) human GGTase-I (yWS3169, *ram2Δ ydj1Δ* [*CEN HIS3 P_PGK1_-FNTA*] [*CEN LEU2 P_PGK1_-PGGT1B*]). Cultures were grown at 25 ℃ or 37 ℃ (denoted with an *), and total cell lysates were prepared from each transformant condition and analyzed by SDS-PAGE and immunoblot using anti-Ydj1 antibody. Data are representative of two biological replicates. um – unmodified; m – modified.

To extend our studies to human GGTase-I, we performed the gel-shift assay using a yeast strain that overexpresses human GGTase-I in the absence of yeast GGTase-I (Hildebrandt, Sarkar et al. 2023). For sequences with higher NGS E-Scores, we observed a fully modified pattern (e.g., CSFL, CHLF, CAFL) or a doublet pattern (e.g., CLIN, CYVM, CWIT, CYIY, CPIQ), while sequences with lower scores (CETT, CDGE, CASQ, CNTH, CVCG) did not display a mobility shift (**Figure 5D**). This ‘humanized’ GGTase-I (*Hs*GGTase-I) expressing strain was better at modifying non-canonical CXXX motifs relative to endogenous yeast GGTase-I. This could be due to a higher amount of the *Hs*GGTase-I enzyme produced in cells under the constitutive *PGK1* promoter. But, this could also indicate specificity differences between the two species of GGTase-I. Interestingly, mammalian and yeast GGTase-I structures differ in active site architecture and the presence of a non-essential proline-rich region in the mammalian enzyme that is proposed to have regulatory properties but whose functional importance has not yet been resolved (Hagemann, Tasillo et al. 2022). We speculate that these features may contribute to mammalian GGTase-I having distinct specificity relative to yeast GGTase-I in our system.

### Rho1 is an effective reporter for yeast GGTase-I activity

Ydj1 is a naturally farnesylated protein that was adapted to use as a GGTase-I reporter in this study. To determine whether the GGTase-I specificity observed in the context of Ydj1 was also apparent in the context of a naturally geranylgeranylated protein, we extended our studies of GGTase-I specificity using Rho1, a well-characterized canonically modified geranylgeranylated yeast protein. To our knowledge, there are no shunted geranylgeranylated yeast proteins, which would have been the preferred starting point for these studies.

*RHO1* is essential, and canonical modification of Rho1 (CVLL) is required for its function (Ohya, Qadota et al. 1993, Yamochi, Tanaka et al. 1994). Hence, we took advantage of an established Rho1 functional assay to examine the impact of different CXXX sequences on Rho1 activity (i.e., plasmid-loss assay) (**Figure 6A**). In this assay, *rho1Δ* yeast complemented by a *URA3*-marked plasmid encoding wildtype Rho1 (i.e., *rho1*1 [*URA3 RHO1*]) are transformed with a *LEU2-*marked plasmid encoding a Rho1-CXXX variant. Negative selection is applied (i.e., 5-FOA) to recover yeast having lost the *RHO1 URA3*-marked plasmid. In this experimental set-up, yeast can only survive negative selection if they retain a functional Rho1-CXXX variant on the remaining *LEU2*-marked plasmid. To eliminate the possibility of farnesylation occurring to Rho1 and interfering with our results, we used as a starting point a strain that also lacked FTase activity (i.e., *ram1*1 *rho1*1 [*URA3 RHO1*]). To confirm the utility of the plasmid-loss assay for studies of Rho1-CXXX variants, wildtype Rho1(CVLL) was evaluated along with an empty vector. As expected, yeast harboring the *CEN LEU2* plasmid encoding Rho1 (CVLL) supported robust growth after 5-FOA selection whereas yeast harboring an empty vector did not grow (**Figure 6B**). The same results were obtained using the HA-tagged version of wildtype Rho1 (CVLL), while unmodified HA-tagged Rho1-SVLL did not grow.

**Figure 6.**
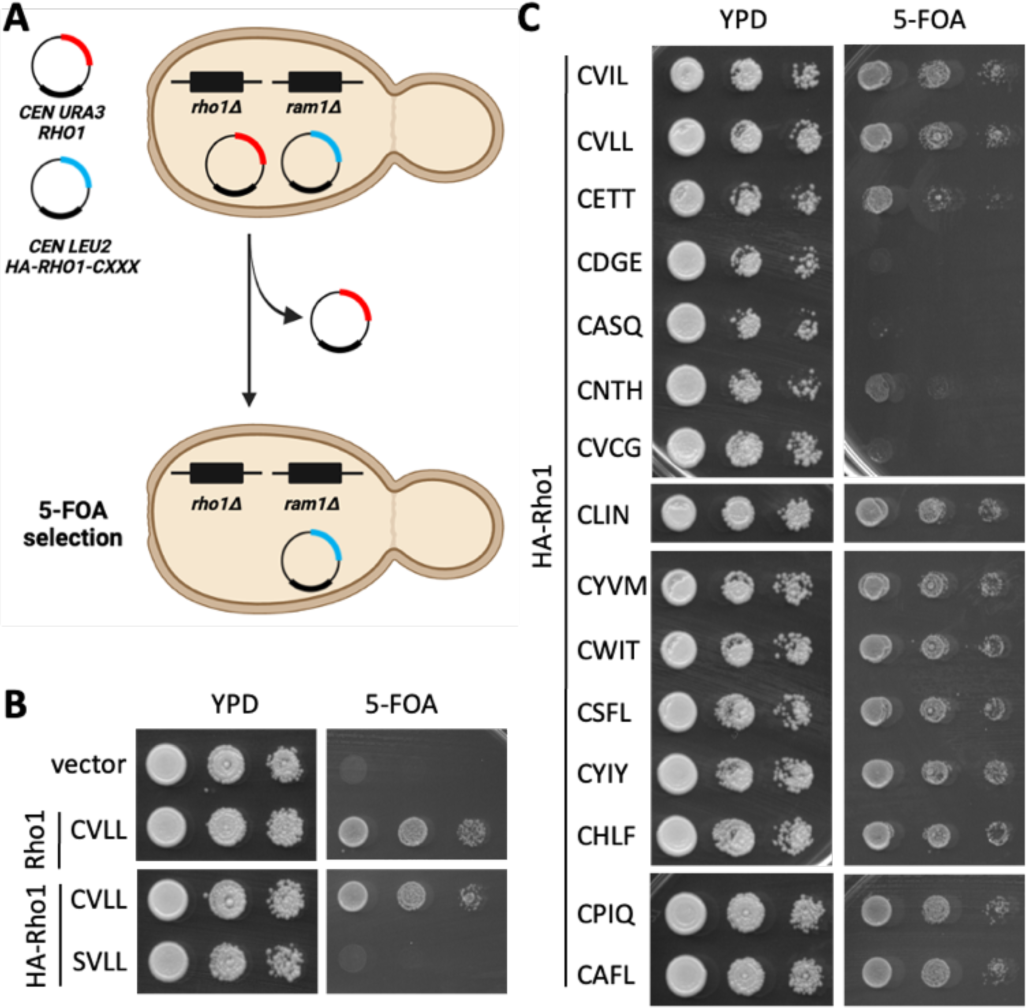
A cell viability assay can distinguish between geranylgeranylated and unmodified Rho1-CXXX variants. **A**) Basis for the plasmid loss assay used to assess the function of Rho1-CXXX variants. The yeast strain has chromosomal-disruptions for the FTase β subunit and Rho1 but is viable due to complementation by a *URA3*-marked plasmid encoding wildtype Rho1 (yWS3761; *ram1*Δ *rho1* [*CEN URA3 RHO1*]). A second *LEU2*-marked plasmid encoding a Rho1-CXXX variant is also present (i.e., *CEN LEU2 RHO1-CXXX*). Upon counterselection with 5-FOA, the *URA3*-marked plasmid is lost, and yeast will only survive counterselection if the *LEU2*-marked plasmid encodes a functional Rho1-CXXX variant. Graphic created using BioRender.com and PowerPoint. **B**-**C)** Yeast transformed with the indicated *CEN LEU2 RHO1-CXXX* plasmids were cultured to saturation in SC-Leucine liquid media, and saturated cultures spotted as 10-fold serial dilutions onto 5-FOA and YPD plates. Similar growth patterns on YPD indicate that the serial dilutions were prepared similarly, while growth on 5-FOA indicates the presence of a functional Rho1-CXXX variant. The 15 candidates in panel C are arranged by increasing NGS E-Score (top to bottom).

To assess the impact of other CXXX sequences on Rho1 function, we applied the plasmid-loss assay to Rho1-CXXX variants designed on the 15 CXXX sequences previously evaluated by Ydj1-dependent thermotolerance and gel-shift assays (**Figure 6C**). This analysis revealed a range of growth patterns for the Rho1-CXXX variants. Those with higher NGS E-Scores (i.e., CLIN, CYVM, CWIT, CSFL, CYIY, CHLF, CPIQ and CAFL) generally grew better than those with lower NGS E-Scores (i.e., CETT, CDGE, CASQ, CNTH and CVCG). Many of the sequences that supported Rho1-dependent growth were non-canonical. Notable exceptions were CVIL and CVLL, which had the two lowest scores within the test set and are expected to be canonically modified geranylgeranylated sequences. While these two sequences underperformed in the context of the Ydj1-dependent thermotolerance assay, they supported Rho1-dependent growth. Overall, the results obtained with the plasmid-loss assay are consistent with the conclusions that geranylgeranylation coupled with canonical modifications hinders Ydj1 but not Rho1 function, and that non-canonical sequences can support Rho1 function.

Considering the results from the Ydj1 and Rho1-based assays together, we predicted that CXXX sequences that poorly supported growth in the context of both reporters (i.e., CETT, CDGE, CASQ, CNTH and CVCG) were not modified by GGTase-I. By contrast, CXXX sequences that supported growth in both contexts (i.e., CYVM, CWIT, CSFL, CYIY, CHLF, CPIQ and CAFL) were predicted to be modified by GGTase-I. These observations can be extended to conclude that the Rho1 function in the context of the plasmid-loss assay is indifferent to canonical vs. likely shunted CXXX modification. An exception to the above binning is the CLIN sequence that did not support Ydj1-dependent thermotolerance but did support Rho1-dependent growth. In this instance, we predict that CLIN is yielding a mixed population of canonically modified and unmodified products, leading to moderate toxicity and a partial yeast growth defect in the context of Ydj1 while the modified population of Rho1 is sufficient to support growth.

### Varied CXXX sequences can support Rho1-dependent growth

Since the function of Rho1 in the plasmid-loss assay did not seem to be impacted by sequences likely to be shunted, we hypothesized that the assay could be adapted into a Rho1-based genetic selection to identify CXXX sequences that can support Rho1-dependent growth, ostensibly because of geranylgeranylation. Importantly, such a Rho1-based screen would be predicted to recover canonically modified CXXX sequences that were not enriched by the Ydj1-based screen.

To test the utility of the plasmid-loss assay for recovering novel functional Rho1-CXXX variants, a random library of plasmid-encoded Rho1-CXXX variants (*LEU2* marked) was expressed in *ram1*1 *rho1*1 [*URA3 RHO1*]) yeast, and the transformed yeast subject to various selections (**Figure 7A**). The library was generated by recombining a PCR product encoding random CXXX sequences into a *LEU2*-marked plasmid encoding a non-functional Rho1 sequence that was designed to lack the entirety of the natural CaaX sequence CVLL (i.e., Rho1-BamHI). While this strategy theoretically allows for the creation of all 8000 possible Rho1-CXXX combinations, only a limited number of plasmid candidates were evaluated due to the labor and time costs of evaluating all of CXXX space. Positive selection (i.e., SC-Leucine) was used to recover colonies harboring recombinant plasmid products, and negative selection of the same colonies (i.e., 5-FOA) was used to identify colonies carrying a functional Rho1-CXXX variant.

**Figure 7.**
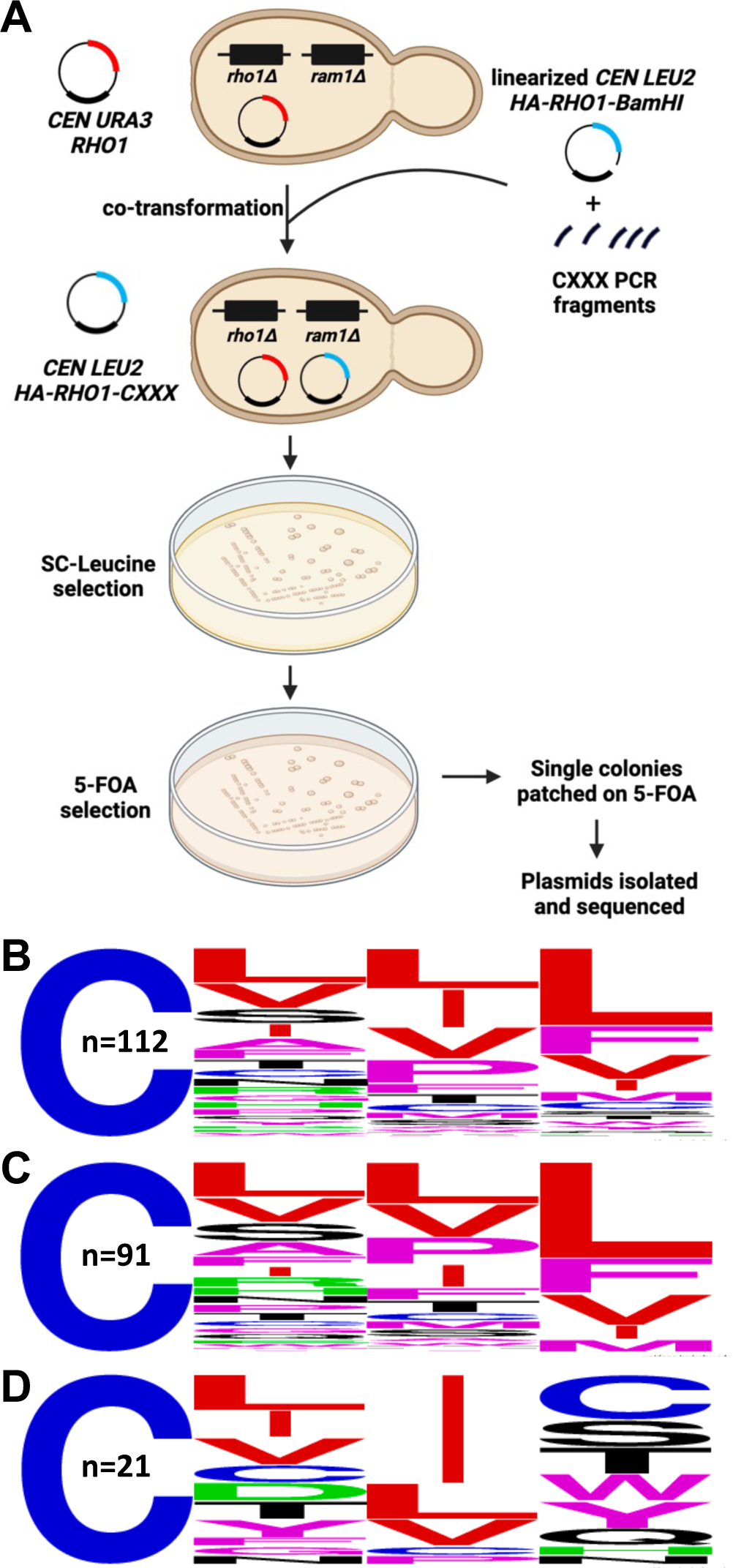
Functional Rho1-CXXX variants can be recovered by the plasmid-loss assay. **A)** Experimental strategy for identifying functional Rho1-CXXX variants. A *ram1*Δ *rho1*Δ [*CEN URA3 RHO1*] yeast background was used for co-introduction of a linearized *LEU2*-based plasmid (*CEN LEU2-HA-RHO1-BamHI*) and PCR products encoding a library of *RHO1-CXXX* sequences. Yeast surviving SC-Leucine selection were replica plated onto 5-FOA media, and plasmids recovered and sequenced from 200 yeast colonies surviving counterselection. Graphic created using BioRender.com and PowerPoint. **B-D**) WebLogo analysis was performed for B) all 112 unique DNA sequences, and sequences conforming to the C) CXX[L/F/I/M/V] consensus and D) CXX[not L/I/F/M/V] consensus.

The Rho1-based genetic screen yielded many 5-FOA resistant colonies. Plasmids were extracted from 200 FOA-resistant colonies and sequenced. Within this set of plasmids, 83 *LEU2*-marked plasmids had the exact DNA sequence of wildtype Rho1 (CVLL) that was encoded in the lost *URA3* plasmid, indicative that gene conversion between the *LEU2* and *URA3* plasmids had likely occurred. Of the remaining 117 sequences, 5 were exact duplicate DNA sequences of other sequences within the set, indicative that their parent colonies were likely double picked during the screening process. Among the remaining 112 unique DNA sequences, some encoded for the same CXXX sequence through different codon usage. In all, 94 distinct CXXX sequences were encoded by the 112 unique DNA sequences, including CSFL and CVIL that are naturally associated with geranylgeranylated proteins (*Hs*Gγ5 and *Sc*Rho5). The 117 hits not matching the parent plasmid DNA sequence were retested using the cell viability assay, and all supported growth on 5-FOA, with most supporting growth that was indistinguishable from that supported by the CVLL sequence (**Figure S4**).

Overall, the Rho1-based screen retrieved CXXX sequences best categorized as a mix of canonical and non-canonical CaaX sequences. WebLogo analysis of the 112 unique hits revealed a prevalence of branched-chain aliphatic amino acids (i.e., L/I/V) at both a_1_ and a_2_ positions (more so at a_2_) and L/F/I/M/V at the X position (**Figure 7B**). These results are consistent with the Caa[L/F/I/M/V] consensus motif reported for geranylgeranylated sequences. Yet, there were clearly sequences recovered from the Rho1-CXXX screen that do not fully match the consensus. About 80% of the hits with a consensus residue at the X position (i.e., L/F/I/M/V) did not have branched-chain aliphatic amino acids at a_1_ or a_2_, indicating some flexibility at these positions (**Figure 7C**). For the approximately 20% of the hits not having a consensus amino acid at the X position (i.e., not L/F/I/M/V), aliphatic amino acids were more strongly prevalent at a_2_ (**Figure 7D**).

A predicted outcome of the Rho1-based screen was that it would recover both canonical and non-canonical CXXX sequences. Moreover, we expected that non-canonical sequences recovered in the Rho1-based screen would be among those enriched in the Ydj1-based screen, while canonical sequences would be de-enriched. Indeed, this is what was generally observed when the 94 unique CXXX hits from the Rho1-based screen were evaluated in the context of their NGS E-Score plots (**Figure 8A and 8B**). Among the 94 unique sequences, about one-half had an NGS E-Score (42 °C vs. naïve *Sc* library) above 2 (n=44; 47%), indicative of enrichment in the Ydj1-based screen. Most of these sequences (n=38) contained a canonical X residue (i.e., L/F/I/M/V) but had mostly non-canonical a_1_ and a_2_ amino acids (**Figure 8C**). About one-third of the unique sequences had an NGS E-Score less than 0.5 (n=34, 36%), indicative of de-enrichment in the Ydj1-based screen. Most of these sequences (n=27) had a canonical Caa[L/F/I/M/V] profile (**Figure 8D**).

**Figure 8.**
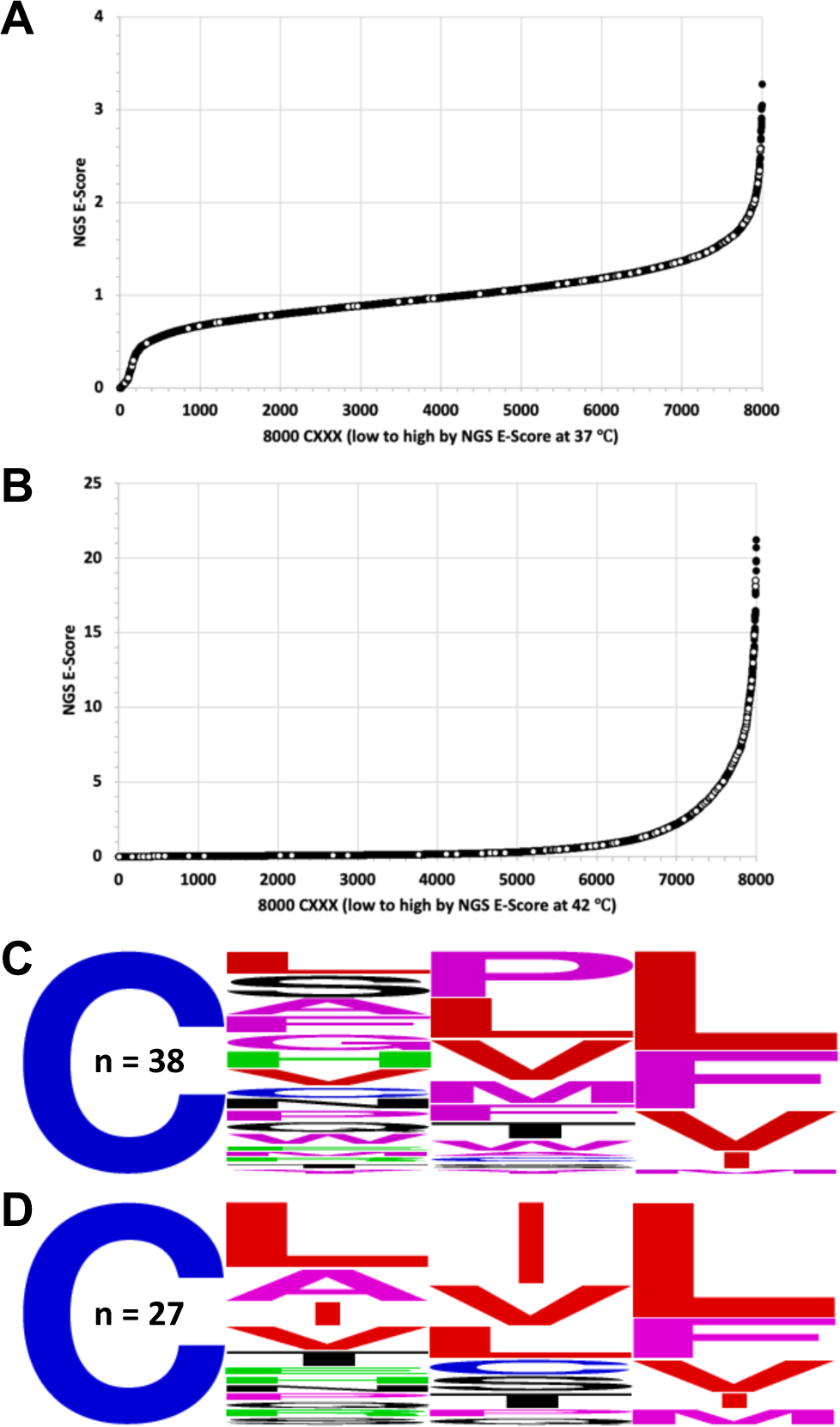
Distribution of NGS-E scores for CXXX sequences identified by Rho1-based screening. **A-B**) The 94 CXXX sequences identified by Rho1-based screening are superimposed as white dots on the NGS E-Score plots (A, 37 ℃ vs. naive yeast library; B, 42 ℃ vs. naive yeast library) derived from the Ydj1-based screen described in **Figures 3A and 3B**. **C-D**). WebLogo analysis of CXXX sequences identified by Rho1-based screening that match the CXX[L/F/I/M/V] consensus and have an NGS E-Score C) >2 or D) <0.5 in the 42 °C data set.

From the set of 94 unique sequences recovered by the Rho1-based screen, four were investigated by Ydj1-based thermotolerance and gel-shift assays (**Figure S5**). These sequences were chosen because they either had high NGS E-Scores (42 °C vs. naïve *Sc* library) in the Ydj1-based assay (i.e., CATL and CNPL) or low NGS E-Scores (i.e., CAIV and CTVL). As expected, Ydj1-CXXX variants encoding CATL and CNPL were able to support thermotolerant growth, while those encoding CAIV and CTVL were not. Also as expected, Ydj1-CXXX variants encoding CAIV and CTVL exhibited a strong gel-shift, while those encoding CATL and CNPL exhibited a very light smear beneath the main Ydj1 band that was absent in the corresponding serine mutants. These results are fully consistent with the observation that all four sequences contain a consensus X residue that is compatible for geranylgeranylation (i.e., L or V), and that the de-enriched sequences have a_2_ residues compatible for cleavage by Rce1 (i.e., I and V), while the enriched sequences do not (i.e., T and P) (Trueblood, Boyartchuk et al. 2000, Berger, Kim et al. 2018, Berger, Yeung et al. 2022).

## Discussion

Since the recognition of prenylation on yeast mating factors and subsequently on mammalian Ras and Ras-related GTPases, the protein prenyltransferases FTase and GGTase-I have been reported to target the “CaaX” consensus motif (Kamiya, Sakurai et al. 1978). Recent studies have revealed, however, that non-canonical CXXX proteins can be farnesylated, but the specificity of geranylgeranylation in this regard has not been fully investigated (Hougland, Hicks et al. 2010, London, Lamphear et al. 2011, Hildebrandt, Cheng et al. 2016, Berger, Kim et al. 2018, Storck, Morales-Sanfrutos et al. 2019, Kim, Hildebrandt et al. 2023). This work addressed this gap in knowledge through two genetic screens whose results demonstrate that yeast GGTase-I has relatively more focused specificity than yeast FTase.

After probing all of CXXX sequence space with the Ydj1 reporter, we determined that consistent features among the best yeast GGTase-I targets were hydrophobic residues at both a_2_ and X, with a dominance of L/F/I/M/V at the X position (**Figure 3A**, bottom WebLogo; **Figure 5B**). Additionally, there was a strong preference for branched-chain aliphatic amino acids (BCAs) at a_2_, unlike yeast FTase that accommodates a wider range of residues at this position. These preferences for a_2_ and X amino acids are also preserved among the sequences recovered using the Rho1 reporter (**Figure 7B**). We speculate that the a_2_ requirement for BCAs in GGTase-I targeted sequences is a means to ensure coupling to the Rce1 protease step of the canonical CaaX modification pathway, leading to proteins with a highly hydrophobic C-terminus (**Figure 1**). Consistent with this premise, geranylgeranylated proteins are typically associated or predicted to be associated with membranes, whereas farnesylated proteins, especially those that are shunted (e.g., Hsp40s, Nap1, Lkb1/Stk11, etc.), are not necessarily membrane associated.

Our results initially suggested that non-canonical sequences might also be GGTase-I targets (**Figures 3A, 3B**), but orthogonal biochemical validation revealed that these sequences are, at best, weakly modified by yeast GGTase-I (**Figures 5A, 5B**). Nonetheless, it appears that limited modification can impart functional properties to both Ydj1 and Rho1. It remains unclear how many geranylgeranylated proteins require partial modification for their function, but having a non-canonical CaaX sequence may provide a means to this end. It also remains unclear whether upregulation of GGTase-I activity could lead to better modification of otherwise poorly modified CaaX sequences, which was observed when overproducing yeast and mammalian GGTase-I in our system (**Figures 5C, 5D**). In natural biological settings, we speculate that increased GGTase-I production could occur in response to external signaling events (e.g., increased transcription) or regulatory mechanisms (e.g., phosphorylation); the latter has been postulated to modulate mammalian FTase activity (Goalstone, Carel et al. 1997). Lastly, it remains unclear whether any non-canonical CaaX sequences displaying partial modification could be better modified in their natural protein context.

Overall, we observe similar target specificities for yeast and mammalian GGTase-I despite substantial differences in active site residues (**Table S1**). Crystallographic studies have revealed that the a_2_ and X binding pockets of mammalian GGTase-I are hydrophobic in nature (Taylor, Reid et al. 2003, Reid, Terry et al. 2004, Gangopadhyay, Losito et al. 2014). These same studies indicate that mammalian GGTase-I has considerable flexibility at the a_1_ position. Yeast GGTase-I appears to have the same specificity features based on the results of our study. This suggests that the yeast and mammalian enzymes have similar active site architecture despite differences in active site residues. Indeed, the overall architecture of the yeast GGTase-I β subunit (Cdc43), as predicted by structural modeling, has a high degree of structural alignment with the established structure of the mammalian GGTase-I β subunit (**Figure 9**; RMSD = 1.3 Å). A key structural difference appears to be the disposition of W108β in yeast GGTase-I, which would clash sterically with the geranylgeranyl group if it were positioned in the active site as for mammalian GGTase-I. We expect that future structural studies of yeast GGTase-I will resolve how the enzyme accommodates the geranylgeranyl group in a different position.

**Figure 9.**
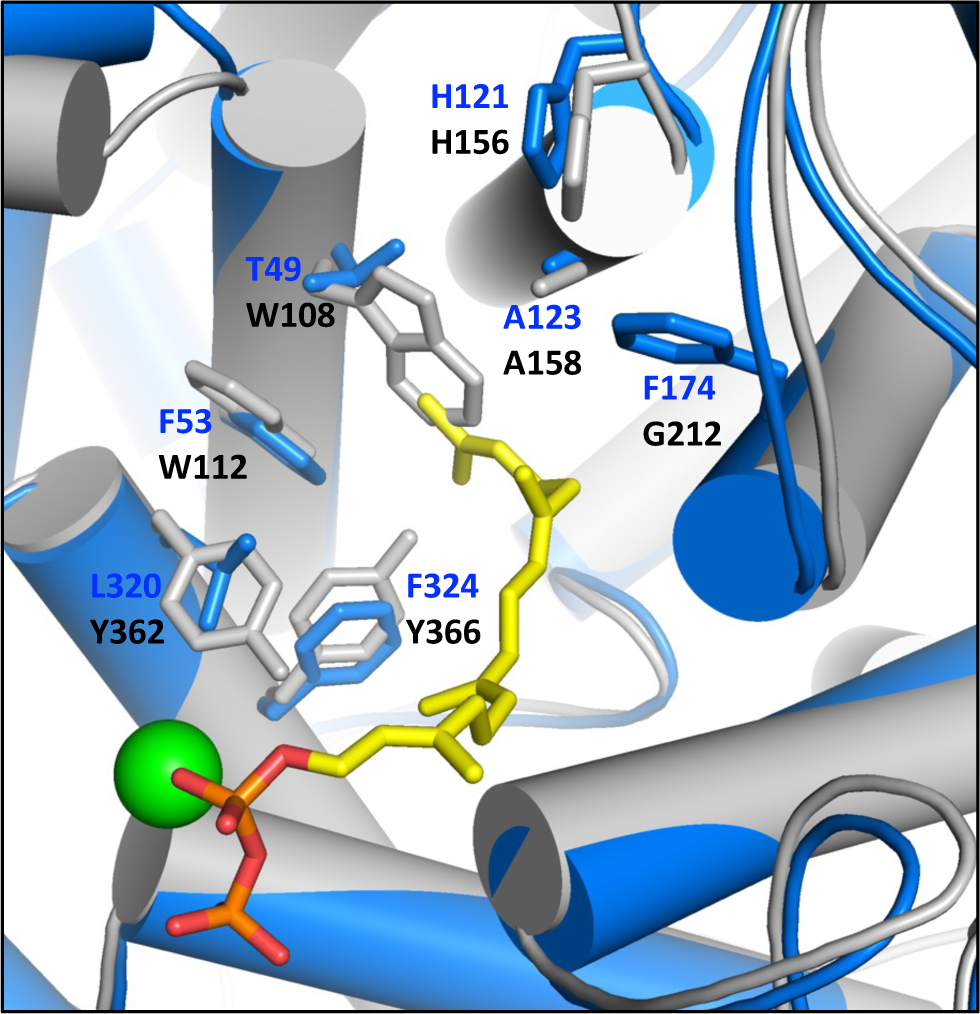
Structure-based alignment of rat and yeast GGTase-I β subunits. The structure of the rat (blue) and yeast (grey) GGTase-I β subunits were derived from PDB 1n4p and an AlphaFold predicted structure, respectively. Active site amino acids are color coded to match the structures; the active site zinc ion is the green sphere; the geranylgeranyl pyrophosphate is indicated in yellow and other colors. Alignment of the structures and an RMSD calculation were performed using the Align function of PyMol.

Our previous studies have revealed that yeast FTase, and likely mammalian FTase, has broader tolerance than previously predicted, especially for a_1_ and a_2_ amino acids. By comparison, our evidence reveals that yeast GGTase-I has a much more focused specificity, consistent with reports for mammalian GGTase-I. Both the yeast prenyltransferases can allow various amino acids at a_1_. While a broader tolerance is exhibited by yeast FTase at a_2_ (i.e., D, E, K, and R are disfavored), GGTase-I appears much more restrictive at this position (i.e., BCAs are favored for well-modified sequences) (Kim, Hildebrandt et al. 2023). Similarly, yeast FTase has broader tolerance at X (i.e., K, P, and R are disfavored) relative to GGTase-I that strongly prefers a limited set of hydrophobic residues at X (i.e., L/F/I/M/V) (Kim, Hildebrandt et al. 2023). Altogether, our comprehensive analysis of yeast GGTase-I specificity reveals that it has a distinct and more restricted target sequence specificity profile than yeast FTase.

## Data availability

The yeast strains and plasmids used in this study will be available upon request. **File S1** contains the CXXX sequence frequencies of the *E. coli*, naïve yeast, 25 °C, 37 °C and 42 °C libraries and the NGS E-Scores. The authors affirm that all data necessary for confirming the conclusions of the article are present within the article, figures, and tables.

## Acknowledgements

We thank Dr. Avrom Caplan (City College of New York) for the anti-Ydj1 antibody, Drs. David Pellman (Dana Farber Cancer Institute, Harvard Medical School) and Satoshi Yoshida (Waseda University) for the SP319 Rho1 plasmid, Dr. James L. Hougland (Syracuse University) for providing valuable feedback on the manuscript, Dr. Zachary A. Wood (University of Georgia) for assistance with Pymol, and Schmidt lab members for constructive feedback and technical assistance.

## Supplementary Tables

**Table S1.** Active site residues of β subunits in rat and yeast GGTase-I.

| $\beta$ subunit | a<sub>2</sub> specificity | X specificity | C20 specificity |
| --- | --- | --- | --- |
| *Rn*PGGT1B<sup>a</sup> | Thr49, Phe53, Leu320 | Thr49, **His121**<sup>c</sup>, **Ala123**, Phe174 | Thr49, Phe324 |
| *Sc*Cdc43<sup>b</sup> | Trp108, Trp112, Tyr362 | Trp108, **His156**, **Ala158**, Gly212 | Trp108, Tyr366 |

<sup>a</sup>Previously reported (Taylor, Reid et al. 2003, Reid, Terry et al. 2004) or <sup>b</sup>determined by structure-based sequence alignment using the Align function of PyMol with the structures of *Rn*PGGT1B (PDB 1n4p) and *Sc*Cdc43 (AlphaFold) (Jumper, Evans et al. 2021). <sup>c</sup>Residues that are conserved based on structure-based sequence alignment are indicated in bold.

**Table S2.** Yeast plasmids used in this study.

| Plasmid number | Genotype | Source |
| --- | --- | --- |
| pRS315 | *CEN LEU2* | (Sikorski and Hieter 1989) |
| pRS316 | *CEN URA3* | (Sikorski and Hieter 1989) |
| pRS413 | *CEN HIS3* | (Sikorski and Hieter 1989) |
| pRS425 | *2μ LEU2* | (Sikorski and Hieter 1989) |
| pWS942 | *CEN URA3 YDJ1* | (Hildebrandt, Cheng et al. 2016) |
| pWS1132 | *CEN URA3 YDJ1-SASQ* | (Hildebrandt, Cheng et al. 2016) |
| pWS1321 | *CEN URA3 YDJ1-CVLL* | (Hildebrandt, Sarkar et al. 2023) |
| pWS1461 | *CEN URA3 YDJ1-CSFL* | (Berger, Yeung et al. 2022) |
| pWS1635 | *CEN URA3 YDJ1-CVIL* | (Hildebrandt, Sarkar et al. 2023) |
| pWS1767 | *CEN URA3 RAM1* | This study |
| pWS1775 | *CEN URA3 YDJ1-CXXX* | (Kim, Hildebrandt et al. 2023) |
| pWS1807 | *2μ LEU2 RAM1* | This study |
| pWS1835 | *CEN URA3 RHO1* | This study |
| pWS1873 | *CEN URA3 YDJ1-SSFL* | This study |
| pWS1885 | *CEN LEU2 P<sub>PGK</sub>-FNTA* | (Hildebrandt, Sarkar et al. 2023) |
| pWS1894 | *CEN LEU2 RHO1* | This study |
| pWS1934 | *CEN LEU2 P<sub>PGK</sub>-PGGT1B* | (Hildebrandt, Sarkar et al. 2023) |
| pWS2101 | *CEN LEU2 HA-RHO1* | This study |
| pWS2102 | *CEN LEU2 HA-RHO1-CSFL* | This study |
| pWS2125 | *CEN LEU2 HA-RHO1-BamH1* | This study |
| pWS2128 | *CEN LEU2 RHO1-SVLL* | This study |
| pWS2129 | *CEN LEU2 HA-RHO1-SVLL* | This study |
| pWS2133 | *CEN URA3 YDJ1-CPLL* | This study |
| pWS2163 | *CEN URA3 YDJ1-CWIT* | This study |
| pWS2164 | *CEN URA3 YDJ1-CNTH* | This study |
| pWS2165 | *CEN URA3 YDJ1-CETT* | This study |
| pWS2167 | *CEN URA3 YDJ1-CDGE* | This study |
| pWS2174 | *CEN URA3 YDJ1-CYVM* | This study |
| pWS2178 | *CEN URA3 YDJ1-CVCG* | This study |
| pWS2197 | *CEN LEU2 HA-RHO1-CASQ* | This study |
| pWS2202 | *CEN URA3 YDJ1-CYIY* | This study |
| pWS2208 | *CEN URA3 YDJ1-CLIN* | This study |
| pWS2225 | *CEN LEU2 HA-RHO1-CVIL* | This study |
| pWS2234 | *CEN URA3 YDJ1-SVLL* | This study |
| pWS2235 | *CEN URA3 YDJ1-SVIL* | This study |
| pWS2237 | *CEN URA3 YDJ1-SNTH* | This study |
| pWS2238 | *CEN URA3 YDJ1-SLIN* | This study |
| pWS2241 | *CEN LEU2 HA-RHO1-CYVM* | This study |
| pWS2243 | *CEN LEU2 HA-RHO1-CWIT* | This study |
| pWS2244 | *CEN LEU2 HA-RHO1-CVCG* | This study |
| pWS2247 | *CEN LEU2 HA-RHO1-CNTH* | This study |
| pWS2249 | *CEN LEU2 HA-RHO1-CLIN* | This study |
| pWS2253 | *CEN LEU2 HA-RHO1-CETT* | This study |
| pWS2254 | *CEN LEU2 HA-RHO1-CDGE* | This study |
| pWS2282 | *CEN URA3 YDJ1-CAPL* | (Berger, Kim et al. 2018) |
| pWS2283 | *CEN URA3 YDJ1-CRPL* | (Berger, Kim et al. 2018) |
| pWS2284 | *CEN URA3 YDJ1-CFAL* | (Berger, Kim et al. 2018) |
| pWS2289 | *CEN URA3 YDJ1-CAFL* | This study |
| pWS2290 | *CEN URA3 YDJ1-CAIV* | This study |
| pWS2291 | *CEN URA3 YDJ1-CATL* | This study |
| pWS2293 | *CEN URA3 YDJ1-CNPL* | This study |
| pWS2294 | *CEN URA3 YDJ1-CPIQ* | This study |
| pWS2297 | *CEN URA3 YDJ1-CTVL* | This study |
| pWS2301 | *CEN URA3 YDJ1-SAFL* | This study |
| pWS2302 | *CEN URA3 YDJ1-SAIV* | This study |
| pWS2303 | *CEN URA3 YDJ1-SATL* | This study |
| pWS2304 | *CEN URA3 YDJ1-SNPL* | This study |
| pWS2305 | *CEN URA3 YDJ1-SPIQ* | This study |
| pWS2306 | *CEN URA3 YDJ1-STVL* | This study |
| pWS2307 | *CEN LEU2 HA-RHO1-CPIQ* | This study |
| pWS2308 | *CEN LEU2 HA-RHO1-CAFL* | This study |
| pWS2309 | *CEN LEU2 HA-RHO1-CHLF* | This study |
| pWS2313 | *CEN URA3 YDJ1-CHLF* | This study |
| pWS2326 | *CEN LEU2 P<sub>PGK</sub>-CDC43* | This study |
| pWS2327 | *CEN LEU2 P<sub>PGK</sub>-RAM2* | This study |
| pWS2350 | *CEN HIS3 P<sub>PGK</sub>-RAM2* | This study |
| SP319 | *CEN URA3 HA-RHO1* | (Yoshida, Bartolini et al. 2009) |

**Table S3:** Plasmid cloning strategies.

| Plasmid number | Construction information | Vector backbone | Insert | Selection media |
| --- | --- | --- | --- | --- |
| pWS1835 | Recombination-mediated PCR-directed plasmid construction in yeast (BY4741) | Restriction enzyme digested linearized pRS316 | PCR product encoding *RHO1* ORF | SC-Uracil |
| pWS1894 | Recombination-mediated PCR-directed plasmid construction in yeast (BY4741) | Restriction enzyme digested linearized pRS315 | *PvuI* digested 1835 | SC-Leucine |
| pWS2101 | Recombination-mediated PCR-directed plasmid construction in yeast (BY4741) | Restriction enzyme digested pWS1894 | *PshAI* and *EcoRI* digested SP319 | SC-Leucine |
| pWS2102 | Recombination-mediated PCR-directed plasmid construction in yeast (BY4741) | Restriction enzyme digested pWS2101 | PCR product encoding C-terminus of *RHO1* with the CXXX sequence altered to CSFL | SC-Leucine |
| pWS2125 | Recombination-mediated PCR-directed plasmid construction in yeast (BY4741) | Restriction enzyme digested pWS2101 | PCR product encoding C-terminus of *RHO1* with CXXX sequence replaced with a *BamHI* site | SC-Leucine |
| pWS2326 | Recombination-mediated PCR-directed plasmid construction in yeast (BY4741) | Restriction enzyme digested pWS1934 | PCR product encoding *CDC43* | SC-Leucine |
| pWS2327 | Recombination-mediated PCR-directed plasmid construction in yeast (BY4741) | Restriction enzyme digested pWS1885 | PCR product encoding *RAM2* | SC-Leucine |
| pWS2350 | Ligation | Restriction enzyme digested pWS2327 | Restriction enzyme digested pRS413 | Ampicillin |

---

**Figure S1.**
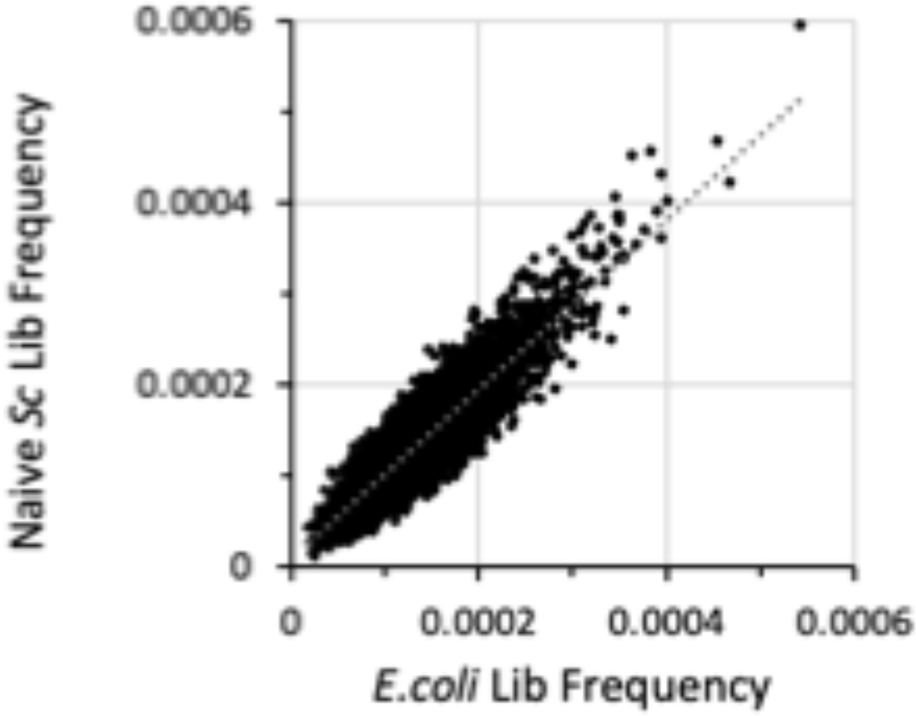
Plot of frequencies observed for YDJ1-CXXX sequences in E. coli and naïve yeast libraries. The frequencies of CXXX sequences within each library are unequal, yielding a range of frequencies. There is, however, a strong correlation (R^2^ = 0.8271) between the frequency distributions observed in the two libraries, indicating no obvious enrichment or de-enrichment for specific YDJ1-CXXX sequences during the yeast transformation process.

**Figure S2.**
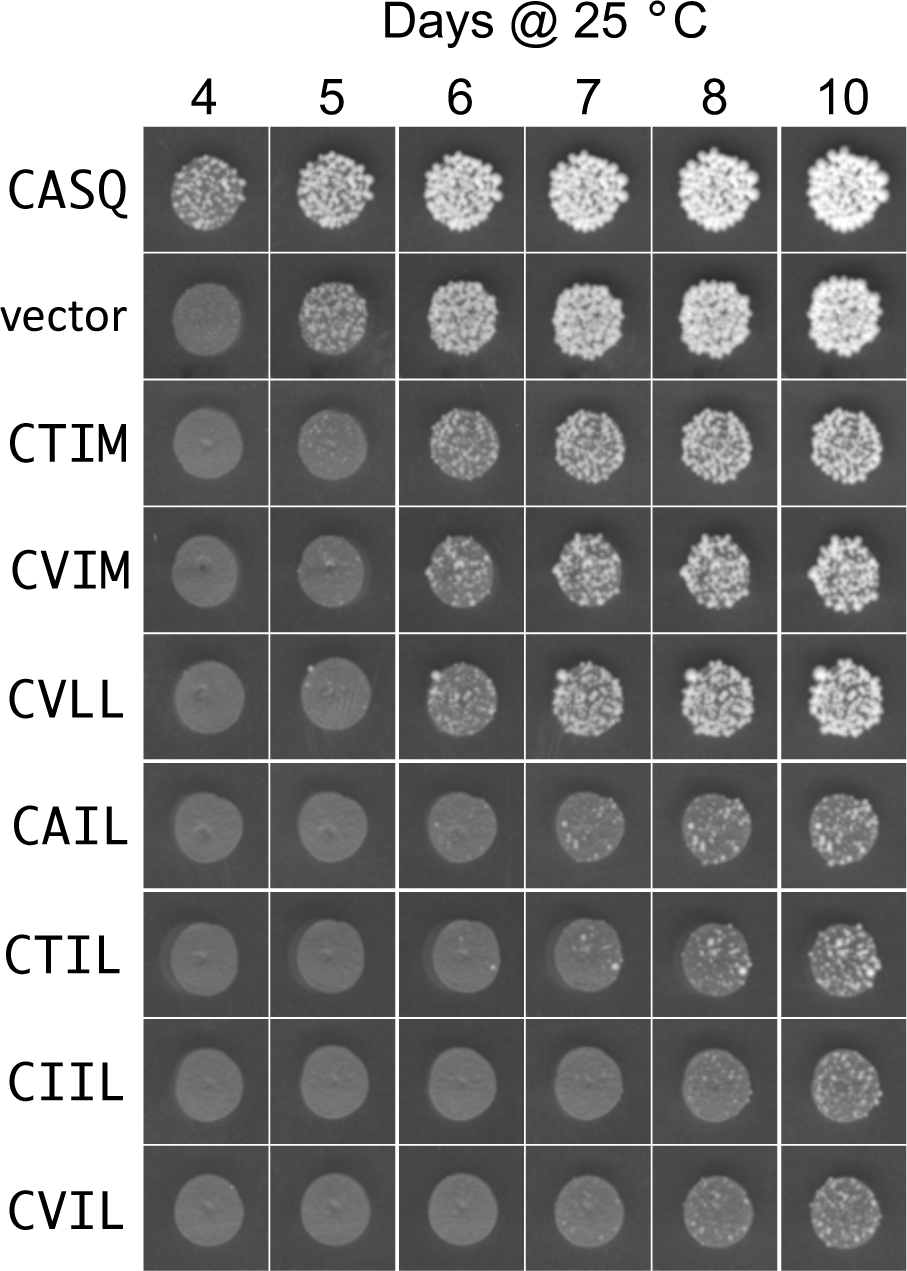
Growth phenotypes of primary transformants expressing Ydj1-CXXX variants with geranylgeranylation potential. Plasmids encoding the indicated Ydj1-CXXX variants were transformed in parallel into yeast lacking *RAM1* and *YDJ1* (yWS2542) using 1 µg of each plasmid. Transformed cells were gently harvested, resuspended to the same volume, and an equivalent portion of each transformation mixture manually spotted onto YPD solid media. Growth of colonies at room temperature was recorded over multiple days.

**Figure S3.**
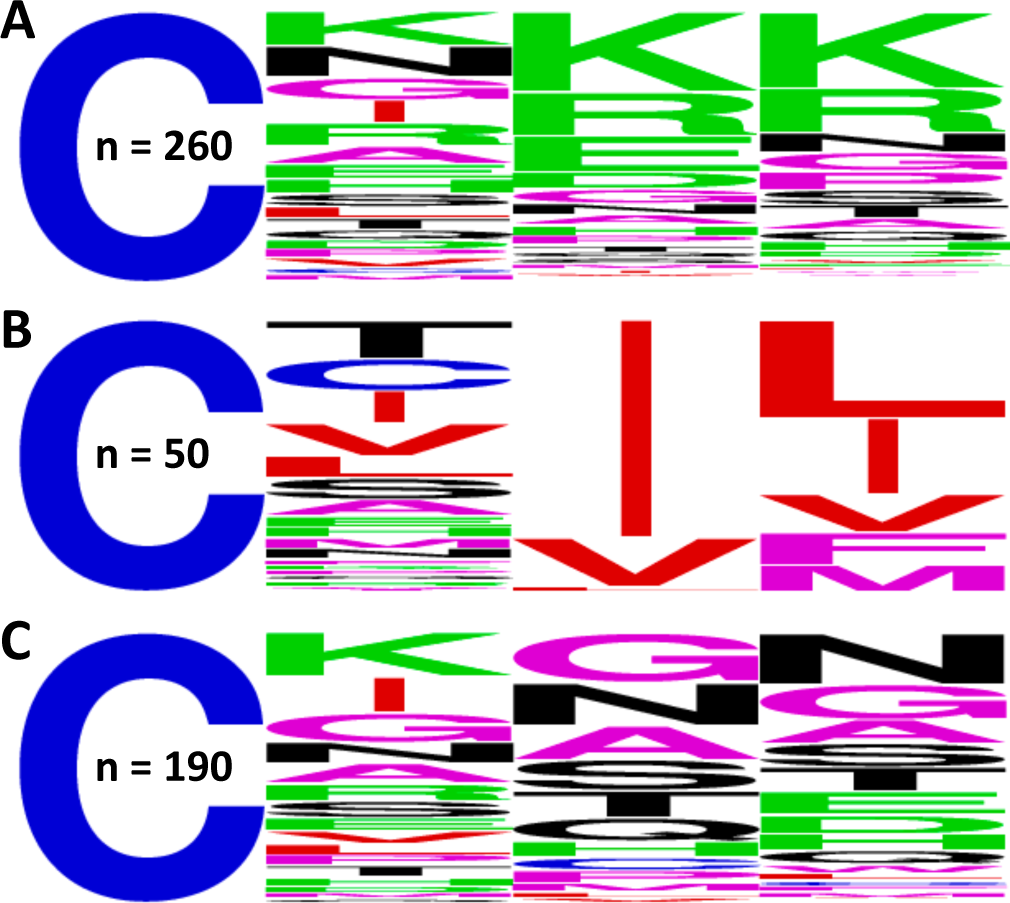
WebLogo analyses of sequence subsets from the lowest 500 NGS E-Scores associated with the 42 °C data set. The sequences evaluated **A**) have D/E/K/R at the a_2_ position or K/P/R at the X position, **B**) match the consensus CX[V/I/L][L/F/I/M/V], or **C**) represent the remaining sequences after removing those evaluated in panels A and B.

**Figure S4.**
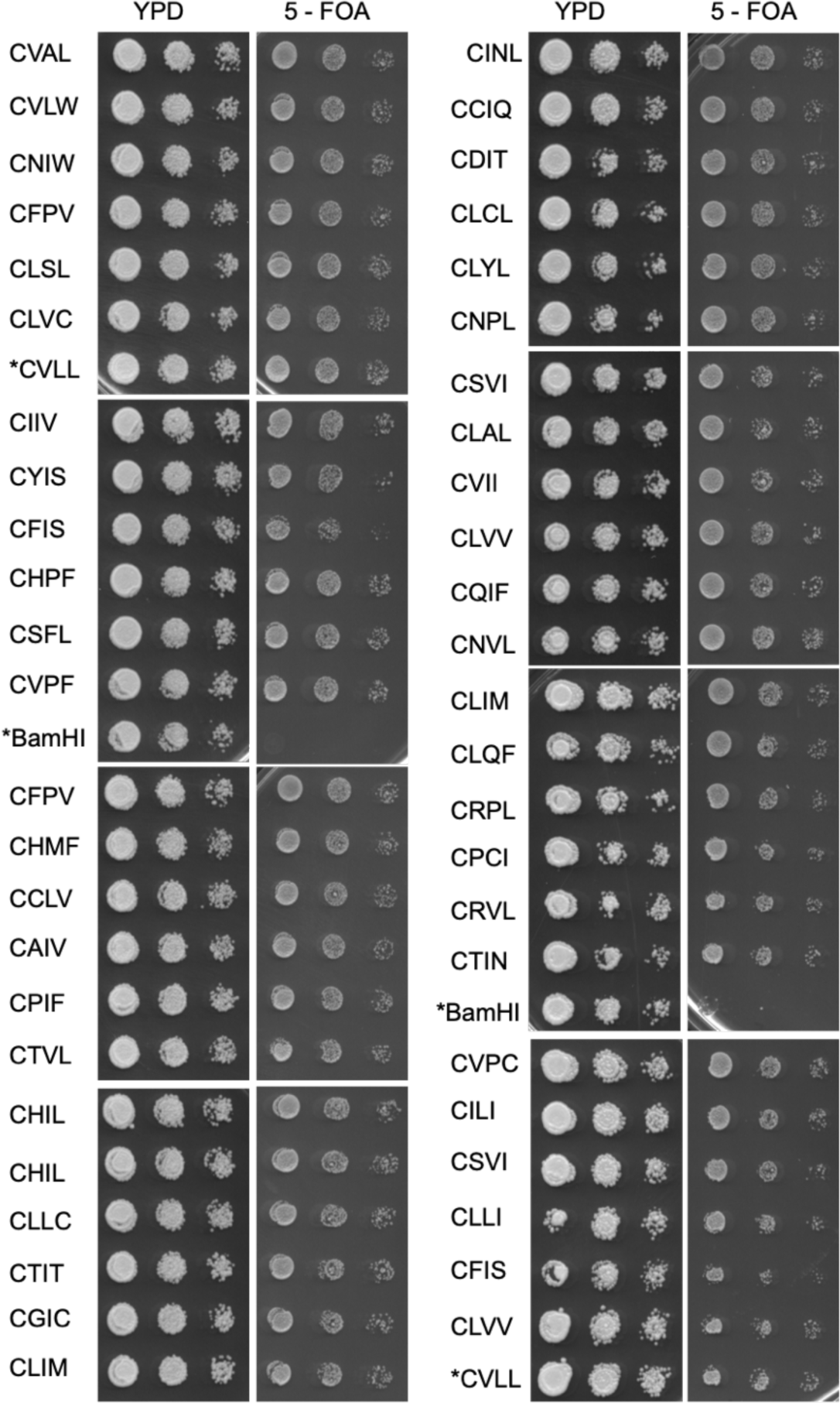
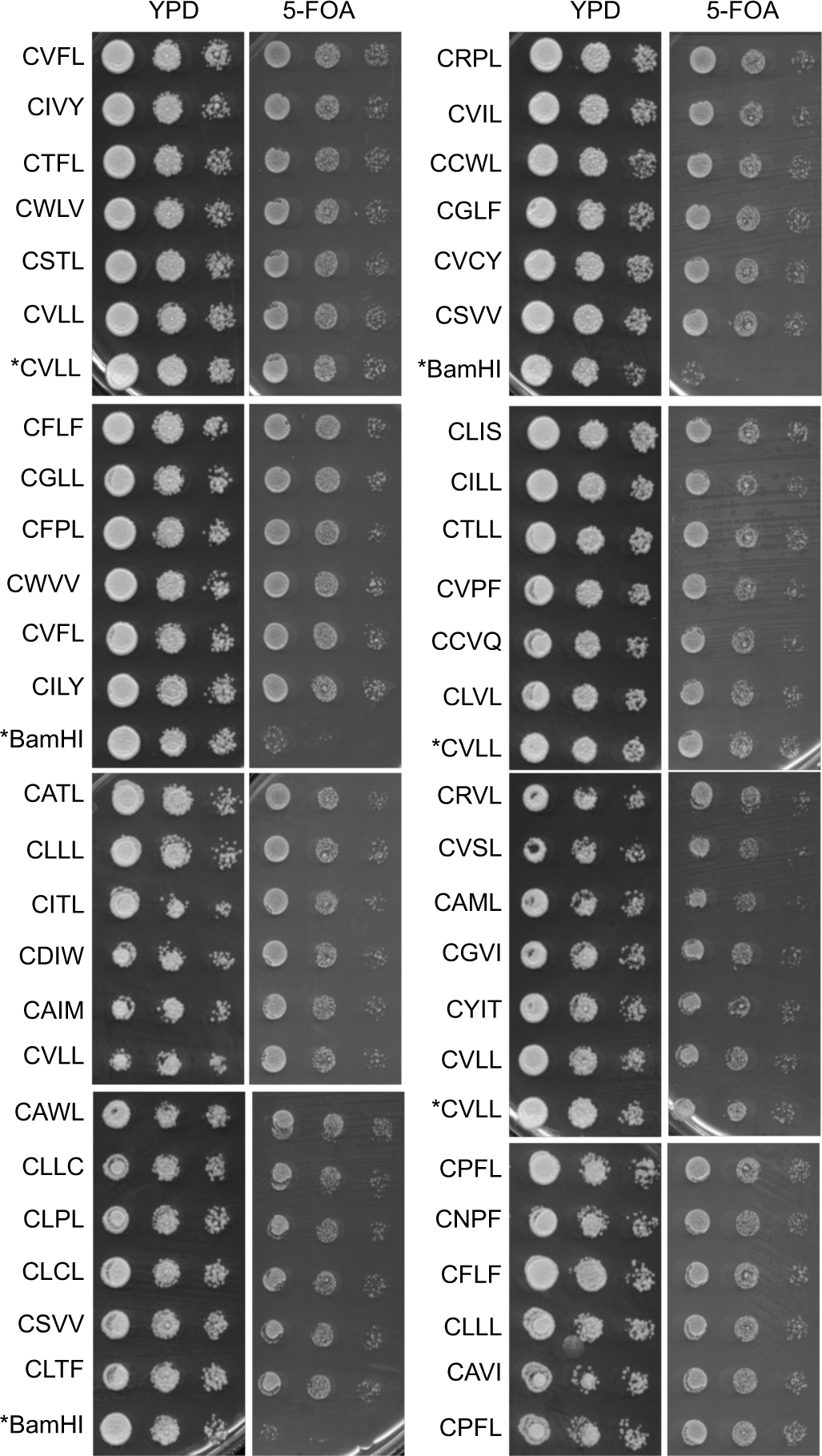
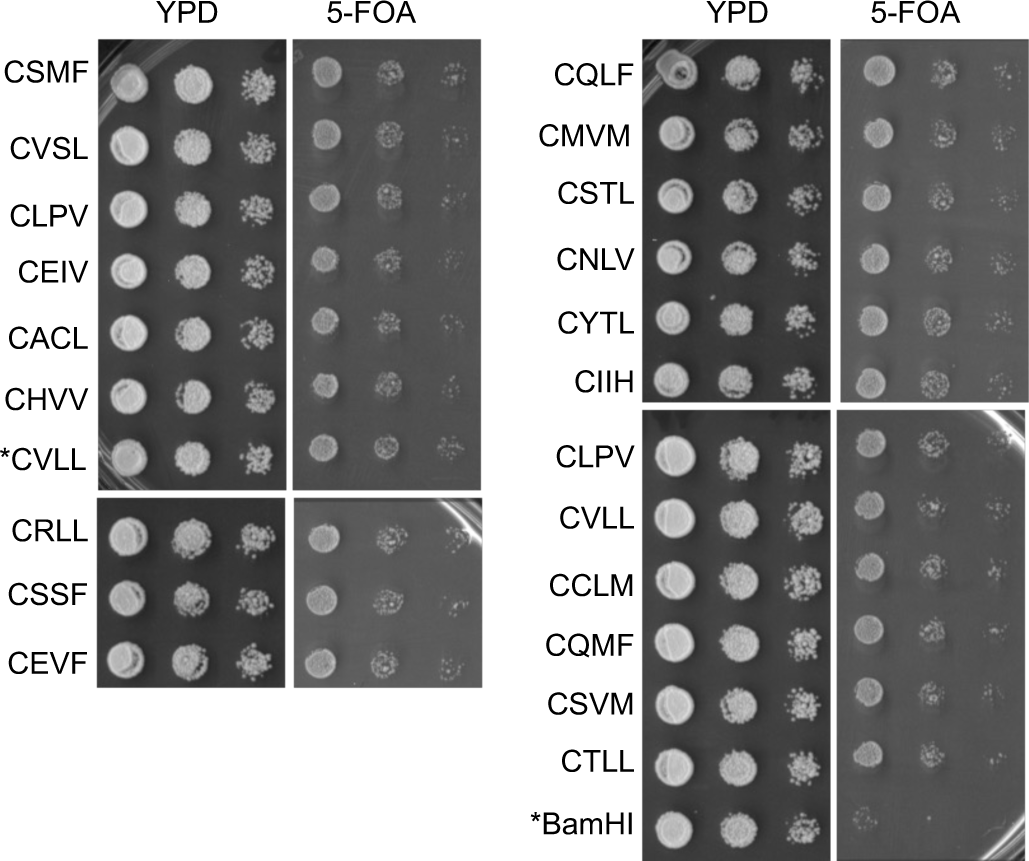
Cell viability phenotypes of Rho1-CXXX variants from Rho1-based screen. The 117 non-parent plasmids recovered by the strategy described in **Figure 7** were transformed individually into yWS3761 (*ram1Δ rho1Δ* [*CEN URA3 RHO1*]) and evaluated by the plasmid-loss assay. For each Rho1-CXXX variant, multiple colonies were used to inoculate SC-Leucine media. Saturated cultures were normalized to 1 A_600_ and used to prepare a 10-fold dilution series that was spotted onto YPD and 5-FOA media. The spotting on 5-FOA media plates was done in two technical replicates. The asterisk (*) denotes transformants expressing wildtype Rho1 (i.e., CVLL) or Rho1 lacking its entire CXXX sequence (i.e., BamHI) that were used as controls and evaluated multiple times.

**Figure S5.**
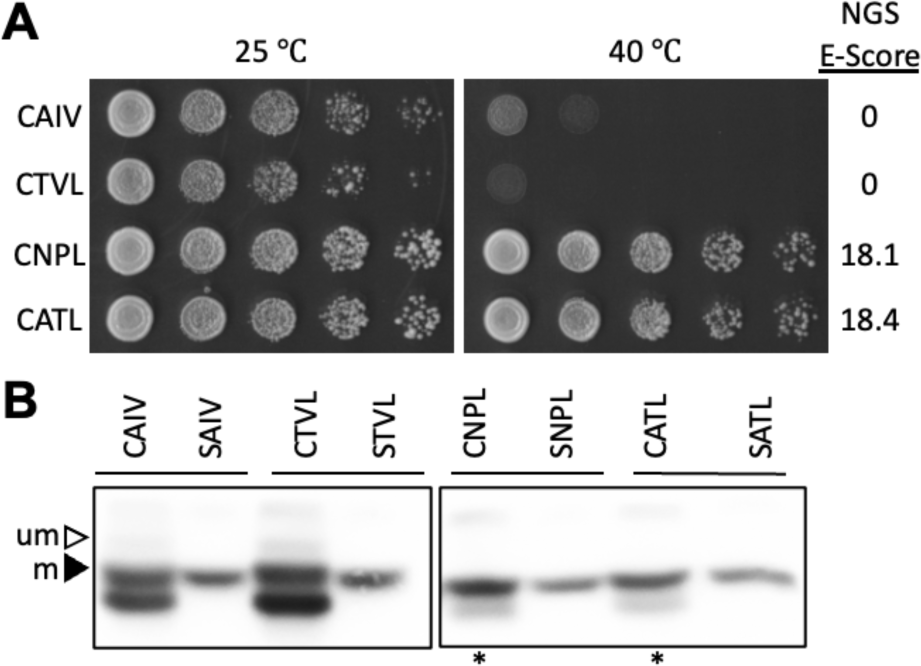
Evaluation of Rho1-based CXXX hits in the context of Ydj1-based assays. Plasmids encoding the indicated Ydj1-CXXX/SXXX variants were evaluated for **A**) thermotolerance and **B**) gel-shift as described in **Figures 1B and 5**, respectively. um – unmodified; m – modified. Data are representative of two biological replicates.

